# Sarm1 is not necessary for activation of neuron-intrinsic growth programs yet required for the Schwann cell repair response and peripheral nerve regeneration

**DOI:** 10.1101/2024.03.04.583374

**Authors:** Ligia B. Schmitd, Hannah Hafner, Ayobami Ward, Elham Asghari Adib, Natalia P. Biscola, Rafi Kohen, Manav Patel, Rachel E. Williamson, Emily Desai, Julianna Bennett, Grace Saxman, Mitre Athaiya, David Wilborn, Jaisha Shumpert, Xiao-Feng Zhao, Riki Kawaguchi, Daniel H. Geschwind, Ahmet Hoke, Peter Shrager, Catherine A. Collins, Leif A. Havton, Ashley L. Kalinski, Roman J. Giger

## Abstract

Upon peripheral nervous system (PNS) injury, severed axons undergo rapid SARM1-dependent Wallerian degeneration (WD). In mammals, the role of SARM1 in PNS regeneration, however, is unknown. Here we demonstrate that *Sarm1* is not required for axotomy induced activation of neuron-intrinsic growth programs and axonal growth into a nerve crush site. However, in the distal nerve, *Sarm1* is necessary for the timely induction of the Schwann cell (SC) repair response, nerve inflammation, myelin clearance, and regeneration of sensory and motor axons. In *Sarm1-/-* mice, regenerated fibers exhibit reduced axon caliber, defective nerve conduction, and recovery of motor function is delayed. The growth hostile environment of *Sarm1-/-* distal nerve tissue was demonstrated by grafting of *Sarm1-/-* nerve into WT recipients. SC lineage tracing in injured WT and *Sarm1-/-* mice revealed morphological differences. In the *Sarm1-/-* distal nerve, the appearance of p75^NTR^+, c-Jun+ SCs is significantly delayed. *Ex vivo*, p75^NTR^ and c-Jun upregulation in *Sarm1-/-* nerves can be rescued by pharmacological inhibition of ErbB kinase. Together, our studies show that *Sarm1* is not necessary for the activation of neuron intrinsic growth programs but in the distal nerve is required for the orchestration of cellular programs that underlie rapid axon extension.

## Introduction

Injury to the adult mammalian peripheral nervous system (PNS) triggers a robust regenerative response in axotomized neurons, however, restoration of motor and sensory function in humans is often incomplete and associated with the development of neuropathic pain (Maita et al. 2023; Scheib and Höke 2013). Compression injuries to the PNS are common and typically cause axonal transection without rupturing the epineurium (Taylor et al. 2008). A compression injury divides the nerve into three major segments: the proximal nerve, the injury site, and the distal nerve. In mice, a sciatic nerve crush injury (SNC) results in limited neuronal death, and the vast majority of severed axons within the proximal nerve persist and initiate regenerative growth. At the injury site, nerve compression causes mechanically induced axon degeneration and unprogrammed (necrotic) death of nerve resident non-neuronal cells. In the distal nerve stump, severed fibers undergo Wallerian degeneration (WD), an active process resulting in rapid axon fragmentation and disintegration of myelin sheaths (Bosch-Queralt, Fledrich, and Stassart 2023; Conforti, Gilley, and Coleman 2014; Gamage et al. 2017; Girouard et al. 2018).

Upon denervation, Schwann cells (SCs) show a stereotypic injury response, characterized by a remarkable degree of cellular plasticity (Stassart, Gomez-Sanchez, and Lloyd 2024). Activation of c-Jun is necessary for the SC injury response and axon regeneration (Arthur-Farraj et al. 2012). The SC repair response is characterized by downregulation of myelin gene products, upregulation of genes associated with immature SCs, neurotrophic factors, and extracellular matrix molecules that promote axon regeneration (Jessen and Mirsky 2016). An important part of the repair response is the breakdown of myelin sheaths into discrete oval-shaped segments and smaller myelin debris. Repair SCs initiate myelin clearance through autophagy of fragmented myelin sheath (Brosius Lutz et al. 2017; Gomez-Sanchez et al. 2015). Infiltration of blood-borne immune cells is important during later stages of myelin removal, nerve debridement, and axonal regeneration (Bastien and Lacroix 2014; Chen, Piao, and Bonaldo 2015; Kalinski et al. 2020; Napoli et al. 2012; Scheib and Höke 2013; Shamash, Reichert, and Rotshenker 2002). Disruption of SC autophagy can delay nerve regeneration and trigger neuropathic pain (Li et al. 2020; Marinelli et al. 2014). Moreover, repair SCs adopt an elongated morphology and align in columns, so-called bands of Büngner, along which axons grow (Arthur-Farraj et al. 2012; Cattin et al. 2015; Clements et al. 2017; Jessen and Mirsky 2019). Importantly, the repair SC is a transient cell state. If axon regeneration is successful, repair SC transition back to myelinating or non-myelinating (Remak) SCs (Bosch-Queralt, Fledrich, and Stassart 2023); if unsuccessful, repair SC undergo atrophy (Ronchi et al. 2017; Scheib and Höke 2013) and release senescence-associated factors that inhibit axonal regeneration (Fuentes-Flores et al. 2023).

In healthy neurons, nicotinamide mononucleotide adenylyltransferase 2 (NMNAT2) is continuously transported into axons to ensure steady levels of axonal nicotinamide adenine dinucleotide (NAD) by converting nicotinamide mononucleotide (NMN) into NAD (Gilley et al. 2013; Gilley and Coleman 2010; Hicks et al. 2012). Axonal NMNAT2 has a high turnover rate and upon injury, levels rapidly decline in the distal axon, resulting in an increase in NMN and simultaneous drop in NAD. The increased NMN:NAD ratio triggers axon self-destruction or WD. Commensurate with this, WD is greatly delayed in the Wallerian degeneration slow (*Wld^s^*) mouse, overexpressing an NMNAT1 variant (Gilley et al. 2013).

A high NMN:NAD ratio causes activation of the metabolic sensor SARM1 (sterile-α and Toll/interleukin 1 receptor [TIR] motif containing protein 1), a master regulator of WD (Figley et al. 2021; Gerdts et al. 2013; Osterloh et al. 2012). SARM1 has NAD hydrolase activity and is regulated by c-Jun N-terminal kinase (JNK) via phosphorylation at residue Ser-548 (Murata et al. 2018). Activation of SARM1 results in a further decrease in axonal NAD and production of cyclic ADP-ribose (cADPR), which is thought to act as a second messenger to increase Ca^2+^. Elevated intra-axonal Ca^2+^ leads to activation of calpains and degradation of neurofilaments (Essuman et al. 2017; Yang et al. 2013).

Nerve injury not only triggers biochemical processes within axons, but also activation of neuron-intrinsic programs in the cell body, critical for axonal regeneration (Mahar and Cavalli 2018; Blesch et al. 2012; Chandran et al. 2016; Weng et al. 2017; Sahoo et al. 2018; Terenzio et al. 2018). Previous work with injured *Wld^S^* animals reported delayed PNS regeneration (Benavides and Alvarez 1998; Bisby and Chen 1990; Brown, Lunn, and Perry 1992; Martin et al. 2010), and reduced conditioning lesion-induced axon outgrowth (Niemi et al. 2013). The role of vertebrate *Sarm1* in PNS regeneration, however, has not yet been examined.

Here we subjected *Sarm1-/-* mice to PNS injury and analyzed nerve regeneration, using a combination of surgical, histological, physiological, behavioral, cell culture, and biochemistry-based approaches. We show that in PNS injured *Sarm1-/-* mice, regeneration of sensory and motor axons is delayed, associated with faulty nerve conduction, and long-lasting behavioral deficits. Mechanistic studies highlight that defective execution of the SC repair response in *Sarm1-/-* mice contributes to regenerative failure.

## Results

### *Sarm1* regulates the caliber of regenerated PNS axons

To investigate the impact of SARM1-dependent WD on peripheral nerve regeneration, we subjected cohorts of adult C57BL/6 wildtype (WT) and *Sarm1-/-* mice to a mid-thigh sciatic nerve crush (SNC) injury (**Figure 1A**). At 49 days post crush (49 dpc), nerve segments distal to the crush site were harvested for plastic embedding. Axial sections were cut and stained with toluidine blue or processed for ultrastructural analyses to quantify the number of regenerated fibers, axon caliber, and myelination (**Figure 1B, S1A**)(Bartmeyer, Biscola, and Havton 2021). In sham-operated WT mice, the average diameter of nerve fibers (3.75 ± 0.08 µm), axons (1.90 ± 0.01 µm), and myelination (g-ratio = 0.50 ± 0.01), were comparable to parallel processed *Sarm1-/-* mice (3.31 ± 0.12 µm, 1.72 ± 0.05 µm, and 0.51 ± 0.01 µm, respectively) (**Figure 1C-E and Figure S1B-D**). In nerve cross sections, the density of fibers per 100 µm^2^ in WT and *Sarm1-/-* sham-operated nerves was similar (2.0 ± 0.2, and 2.5 ± 0.5, respectively) (**Figure S1E**). At 49dpc, robust axon regeneration is observed in injured WT and *Sarm1-/-* mice, however there was a significant shift toward thinner fibers and axon calibers in *Sarm1-/-* mice (**Figure S1B, C**). In WT mice, the average diameter of regenerated axons and fibers (1.58 ± 0.04 µm and 2.94 ± 0.11 µm) was larger when compared to *Sarm1-/-* mice (1.19 ± 0.09 µm and 2.15 ± 0.21 µm) (**Figure 1C, D**). However, the density of regenerated axons in *Sarm1-/-* nerves (2.8 ± 1) was similar to WT (2.4 ± 0.3) at 49 dpc (**Figure S1E**) and remyelination was comparable, as assessed by calculation of the g-ratio for WT fibers (0.53 ± 0.03) and for *Sarm1-/-* fibers (0.55 ± 0.03) (**Figure 1E, S1D**). Thus, *Sarm1* is not required for normal PNS development, but upon nerve injury regulates the caliber of regenerated axons.

**Figure 1:**
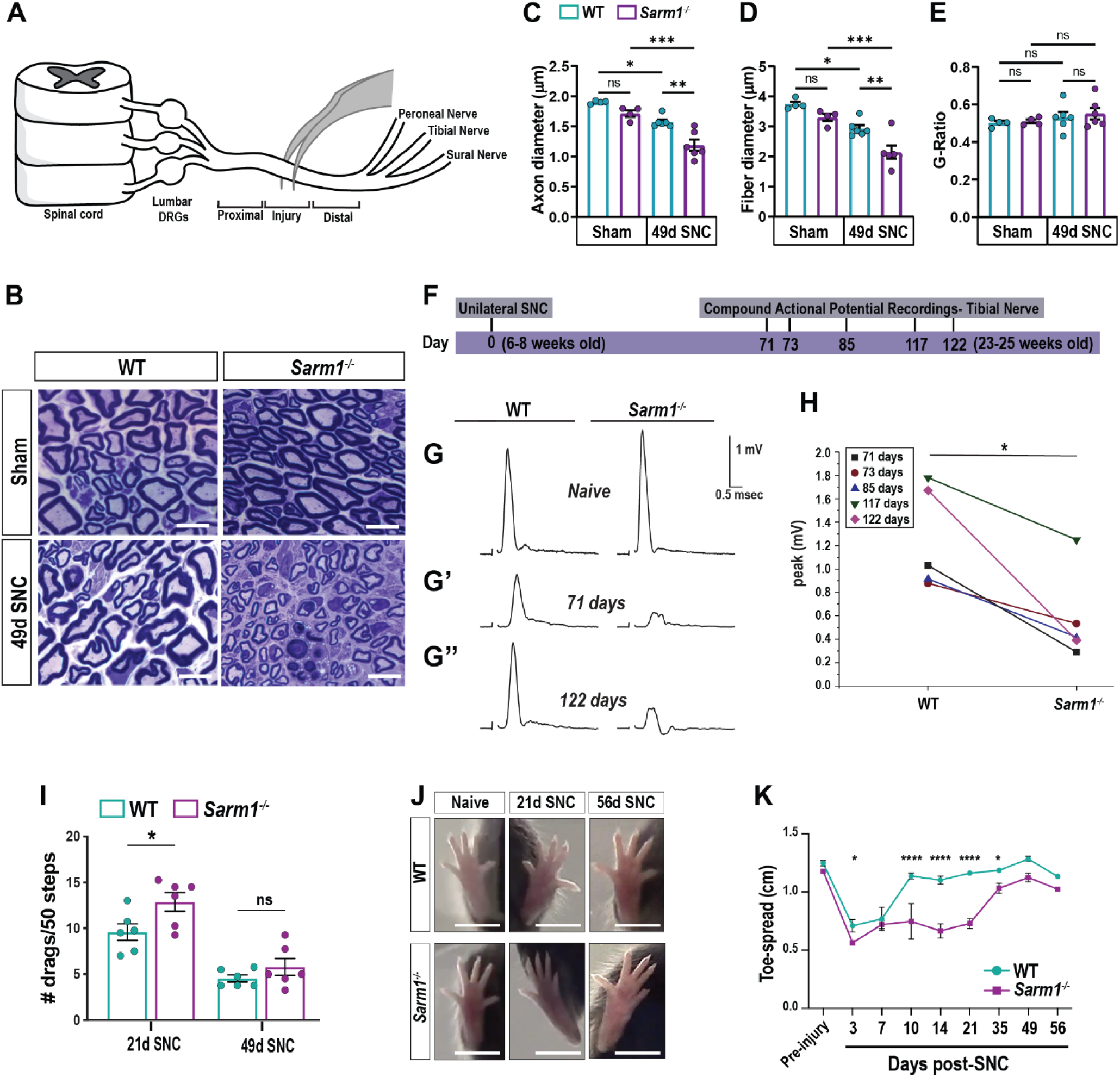
Impaired PNS regeneration in *Sarm1-/-* mice. (**A**) Schematic of mid-thigh sciatic nerve crush (SNC) injury. Brackets demark the location of injury site, proximal, and distal nerve segments. (**B**) Semi-thin sciatic nerve cross sections stained with toluidine blue from sham-operated WT and *Sarm1-/-* mice, and the distal nerve, 49 days post-SNC. Representative images; sham, n = 4 mice per genotype; 49 days post-SNC, n = 6 mice per genotype. Scale bars, 10 µm. (**C-E**) Quantification of axon diameter, fiber diameter, and g-ratio of sham-operated and distal nerve 49 days post-SNC of WT and *Sarm1-/-* mice. Data are represented as mean ± SEM. Student’s t-test with Mann-Whitney post hoc test; *p≤ 0.05; **p≤0.01; ***p≤0.001; ns, not significant. (**F**) Timeline of CAP recordings of acutely excised tibial nerves from naïve and injured WT and *Sarm1-/-* mice. (**G-G’’**) Representative CAP recordings from naïve and injured WT and *Sarm1-/-* mice. Calibrations of amplitude (1 mV) and time (0.5 msec) are shown. (**H**) CAP amplitudes of WT and *Sarm1-/-* tibial nerves post-SNC, recorded after the designated time points. The amplitude in*Sarm1-/-* nerves is significantly reduced. N = 6 injured mice per group, unilateral sciatic nerve crush (left leg; right leg was used as sham control); Student’s t-test; *p=0.014. (**I**) Analysis of foot placements on the horizontal ladder beam. The number of drags per 50 septs is shown for WT (green) and *Sarm1-/-* (purple) mice at 21d and 49d post-SNC. N = 6 mice per genotype, unilateral SNC (left leg). Data are represented as mean ± SEM. Each data point represents one biological replicate. *p = 0.036 by unpaired Student’s t-test. (**J**) Hindfeet, showing toe spreading of sham-operated, 21d, and 56d SNC WT and *Sarm1-/-* mice. Scale bar, 1 cm. (**K**) Longitudinal assessment of toe spreading reflex; mean distance between the tips of the first and fifth toes ± SEM is shown. WT pre-injury, 3d, 21d, and 35d, n = 10 mice; WT 14d and 49d, n = 6 mice; WT 7d, 10d, and 56d, n = 4 mice. *Sarm1-/-* pre-injury, 3d, 21d, and 35d, n = 9 mice; *Sarm1-/-* 14d and 49d, n = 5 mice; WT 7d, 10d, and 56d, n = 4 mice. Two-way ANOVA; *p≤ 0.05; ****p≤ 0.0001.

### PNS injured *Sarm1-/-* mice exhibit lasting deficits in nerve conduction

To assess whether morphological alterations in the regenerated *Sarm1-/-* PNS impact nerve conduction, we measured compound action potentials (CAPs) in the tibial branch of acutely excised nerves (**Figure 1F-H**). Nerve specimens prepared from naïve WT and *Sarm1-/-* mice exhibited comparable conduction velocities (33.6 ± 0.6 m/sec and 30.5 ± 0.6 m/sec) and amplitudes (2.0 ± 0.1 mV and 2.2 ± 0.2 mV), respectively (**Figure 1G**). At 10 dpc, we recorded weak but comparable CAP signals in WT (0.2 ± 0.1 mV) and *Sarm1-/-* (0.3 ± 0.2 mV) nerves (data not shown). To ask whether nerve crush injury to *Sarm1*-/- mice results in long-lasting conduction deficits, CAP recordings were carried out at 71, 73, 85, 117, and 122 dpc. Representative sweeps from 2 time points (71d and 122d) are shown in **Figure 1G’-G’’**, along with sweeps from uncrushed nerves. In injured WT mice, CAP amplitudes significantly increased (p= 0.003) between 71 dpc (mean overall 1.3 ± 0.1 mV) and 122 dpc (2.0 ± 0.1 mV), approaching pre-injury levels. In contrast, CAP amplitudes in injured *Sarm1-/-* mice did not significantly increase (p = 0.25) between 71 dpc (0.6 ± 0.1 mV) and 122 dpc (0.6 ± 0.2 mV), and thus, failed to reach pre-injury levels. This *Sarm1* dependence is evident in **Figure 1H**, where CAP amplitudes of WT and *Sarm1-/-* nerves are plotted as pairs at each of the 5 post-injury time points examined. In every case, the CAP amplitude was higher in WT than in *Sarm1-/-* mice (*p = 0.014). In WT mice, regenerated fiber conduction velocities averaged 19.6 ± 0.8 m/sec, versus 33.6 ± 0.6 m/sec in uncrushed axons (p ≤ 0.001). In *Sarm1-/-* mice, the corresponding values were 16.2 ± 0.3 m/sec and 30.5 ± 0.6 m/sec (p ≤ 0.001). The difference in conduction velocities between WT and *Sarm1-/-* regenerated fibers was not significant. Together, these studies show that PNS regeneration in injured *Sarm1-/-* mice is delayed and functionally incomplete with respect to electrical impulse strength, but not conduction velocity.

### *Sarm1-/-* mice show delayed restoration of hind paw function

To assess functional regeneration, we examined hind paw placement of mice walking across a horizontal ladder (Ribeiro et al. 2017). Naïve WT and *Sarm1-/-* mice were trained and hind paw steps on ladder beams counted and classified as correct steps or foot faults (drags and slips). The same cohort of mice was then subjected to unilateral SNC, and ipsilateral paw placement assessed in a longitudinal study over 49 days. At 3 dpc, hind paw placement was severely impaired with more than 50% of steps scoring as foot faults, independently of genotype (**Figure S1F**). By 21 dpc, hind paw placement in WT mice had improved to 88.3 ± 1.1% success, and was significantly higher than in mutants, scoring 80.6 ± 2.7% success (**Figure S1F**). For normalization, we calculated a ladder beam score (LBS), defined as the number of foot faults on the ipsilateral side per 50 steps (**Figure 1I**). In *Sarm1-/-* mice, at 21 dpc, the LBS was significantly higher than in parallel processed WT mice (**Figure 1I**). At 49 dpc however, the LBS no longer differed between the two genotypes (**Figure 1I**).

As an independent assessment, we compared restoration of the toe-spread reflex on the ipsilateral hind paw of injured WT and *Sarm1-/-* mice (**Figure 1J, K**). Immediately following SNC, the hind paw is clawed. The width of the paw, defined as the distance between the tips of the first and fifth toes, increases as axons regenerate (Wilder-Kofie et al. 2011). Toe-spread prior to SNC was comparable between WT (1.20 ± 0.02 cm) and *Sarm1-/-* mice (1.18 ± 0.02 cm). At 3 dpc, toe-spread reflex was absent, and the toe distance was 0.7 ± 0.05 cm in WT mice and 0.56 ± 0.015 cm in *Sarm1-/-* mice. Between 7-14 dpc, the toe-spread increased from 0.77 cm to 1.1 cm in WT mice, but did not increase (0.72 cm and 0.66 cm) in *Sarm1-/-* mice (**Figure 1J, K**). Between 21-56 dpc, the toe-spread distance remained constant at 1.1 to 1.2 cm in WT mice, and in *Sarm1-/-* mice gradually increased after 21 dpc to reach 1.1 cm at 49 dpc. At 56 dpc, *Sarm1-/-* mice exhibit a toe-spread reflex similar to pre-injury mice. Overall, behavioral studies provide independent evidence that peripheral nerve regeneration is significantly delayed in *Sarm1-/-* mice.

### *Sarm1-/-* mice exhibit delayed motor axon regeneration and aberrant target innervation

Defects in the toe-spread reflex prompted us to assess regeneration and target reinnervation of motor neurons. Specifically, we analyzed the neuromuscular junctions (NMJs) formed by the peroneal branch of the sciatic nerve with muscle fibers in the extensor digitorum longus (EDL), a fast twitch muscle in the lateral part of the lower leg. The pretzel-shaped postsynaptic endplate on individual muscle fibers was labeled with bungarotoxin (BTX), motor axons with anti-class III β-tubulin (Tuj1), and presynaptic terminals with anti-synapsin (Syn) staining. In naïve *Sarm1-/-* mice, NMJs appeared normal, presynaptic motor terminals and postsynaptic sites were in close apposition and there were no obvious differences compared to naïve WT mice (**Figure S2A**). To assess motoneuron axon regeneration and muscle fiber reinnervation, NMJs were analyzed at 7, 14, 21, and 42 dpc. To quantify endplate reinnervation, we followed a previously established 0-5 scoring system (Shadrach and Pierchala 2018). A score of 0 is given for a completely denervated endplate and a score of 5 is given for a fully re-innervated endplate (**Figure S2B**). At 7 dpc, motor axons remain visible at endplates, however staining for the presynaptic marker Syn was greatly reduced, in both WT and *Sarm1-/-* mice (**Figure S2C-E**). Whole mount staining of the EDL muscle revealed axon fragmentation in WT but not *Sarm1-/-* mice (**Figure S2F**). At 14 dpc, WT mice show significantly more axons innervating endplates compared to mutants. Syn+ presynaptic structures are abundantly present at WT NMJs (scores 3-5), but largely missing in *Sarm1-/-* mice (scores 0 and 1) at 14 dpc (**Figure 2A-C**). At 21 dpc, 57% of WT NMJs are fully reinnervated and form mature presynaptic specializations (score 5). In *Sarm1-/-* mice, however, most endplates are incompletely reinnervated (scores of 1 and 2) and less than 1% show full reinnervation (score 5) (**Figure 2D-F**). Similarly, formation of Syn+ presynaptic specialization is lagging in *Sarm1-/-* mice and most NMJs at 42 dpc **(Figure 2G-I)**. Higher magnification images of NMJs at 21 dpc revealed pretzel-like structures in WT mice and more irregular structures in *Sarm1-/-* mice (**Figure S2G**), indicating that *Sarm1-/-* mice exhibit defects in apposition of presynaptic motor terminals and postsynaptic endplates. Together these studies show that *Sarm1* is necessary for timely motor axon regeneration and proper formation of presynaptic specialization.

**Figure 2:**
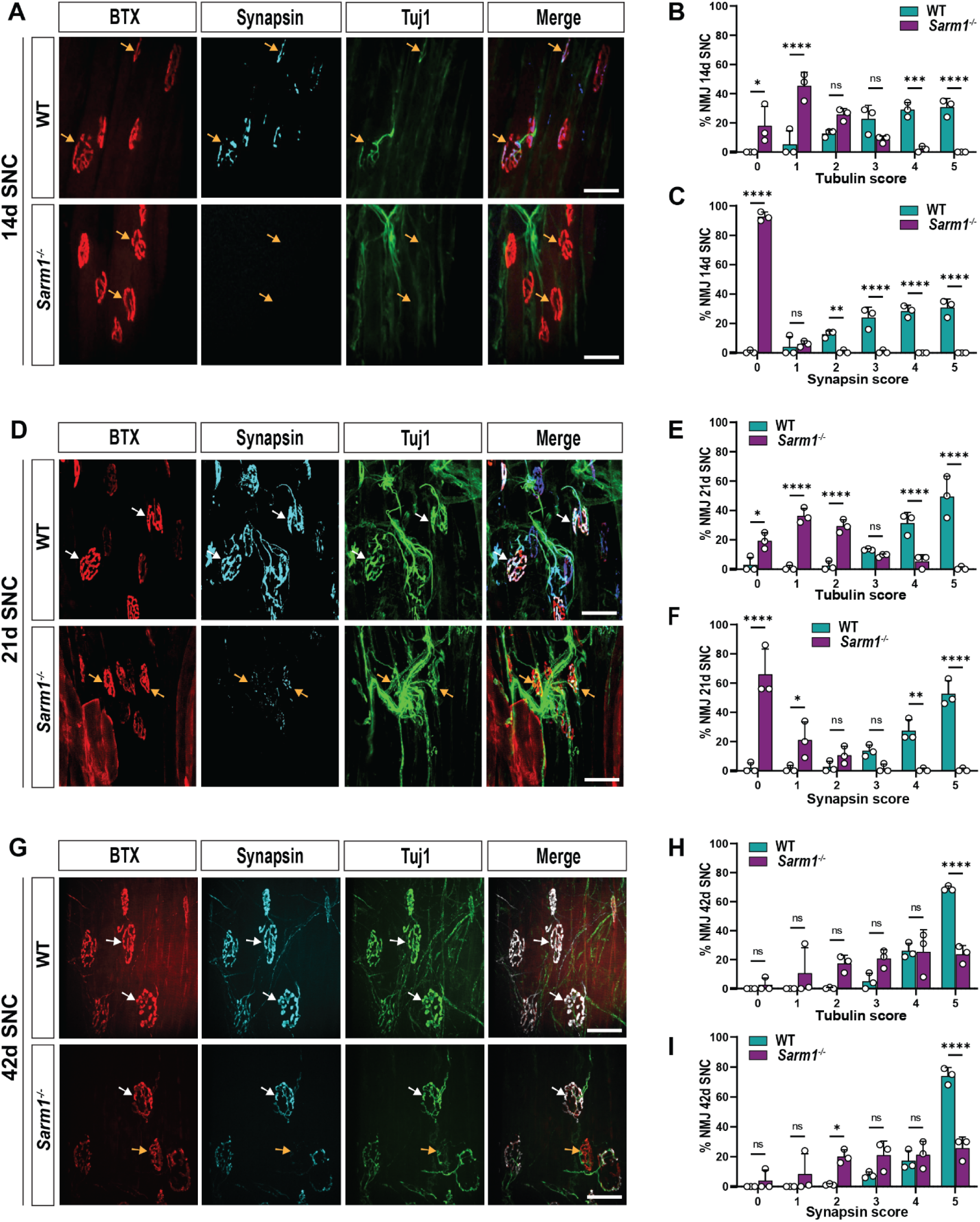
*Sarm1-/-* mice exhibit delayed motor axon regeneration and aberrant endplate innervation. (**A, D**, and **G**) Representative images of whole-mount extensor digitorum longus (EDL) muscles of WT and *Sarm1-/-* at 14, 21, and 42 days post-SNC. Neuromuscular junctions (NMJ) stained with Bungarotoxin, BTX (postsynaptic), synapsin (pre-synaptic), and βIII tubulin; Tuj1 (axon). Yellow arrows indicate incomplete reinnervation (scores 0-4) and white arrows indicate complete reinnervation (score 5). Scale bar, 50 µm. (**B, E,** and **H**) Quantification of NMJ reinnervation by βIII tubulin-labeled axons at 14, 21, and 42 days post-SNC. (**C, F,** and **I**) Quantification of NMJ reinnervation by synapsin-labeled presynaptic terminals at 14, 21, and 42 days post-SNC. Number of NMJ analyzed at 14d (WT = 120; *Sarm1-/-* = 151; n = 3 mice per genotype), 21d (WT = 176, *Sarm1-/-* = 218; n= 3 mice per genotype), and 42d (WT = 221, *Sarm1-/-* = 278; n = 3 mice per genotype). Data are represented as mean ± SEM. *p≤0.05; **p≤0.01; ***p≤0.001, by two-Way ANOVA. Quantification of NMJ innervation at different post-injury time points followed the scoring system where score 0 = fully de-innervated and score 5 = fully innervated (Figure S2 for examples).

### *Sarm1* is required for the timely regeneration of sensory axons in the injured sciatic nerve

To examine sensory axon regeneration following sciatic nerve crush, SCG10 (Stathmin-2)+ axons in longitudinal nerve sections were quantified from WT and *Sarm1-/-* mice. At 3 dpc, sensory axons in WT mice traversed the injury site and entered the distal nerve stump in large numbers (**Figure 3A**). In *Sarm1-/-* mice, sensory axons regenerated robustly into the injury site, but far fewer axons were found in the distal nerve (**Figure 3A**). At 7 dpc, sensory axons in *Sarm1-/-* mice had entered the distal nerve stump; however, in significantly fewer numbers than in 7dpc WT mice (**Figure 3B, B’**). For independent validation of reduced sensory axon regeneration, we collected nerves from injured WT and *Sarm1-/-* mice at 3 and 7 dpc. Nerves were divided into smaller segments (∼3 mm in length) containing proximal nerve, the injury site, or distal nerve (**Figure 3C**). Western blot analysis of WT nerve segments identified SARM1 protein in all segments at 3 and 7 dpc, but not in mutants (**Figure 3D**). Importantly, SCG10 is abundantly present in the proximal nerve and the injury site of both WT and *Sarm1-/-* nerves. In the distal nerve, however, only WT mice show a strong SCG10 signal, while in *Sarm1-/-* distal nerve, SCG10 is significantly reduced at 3 and 7 dpc (**Figure 3D-F**). Together with histological studies, Western blotting shows that injured sensory axons in *Sarm1-/-* mice extend robustly into the injury site but fail to thrive in the distal nerve.

**Figure 3:**
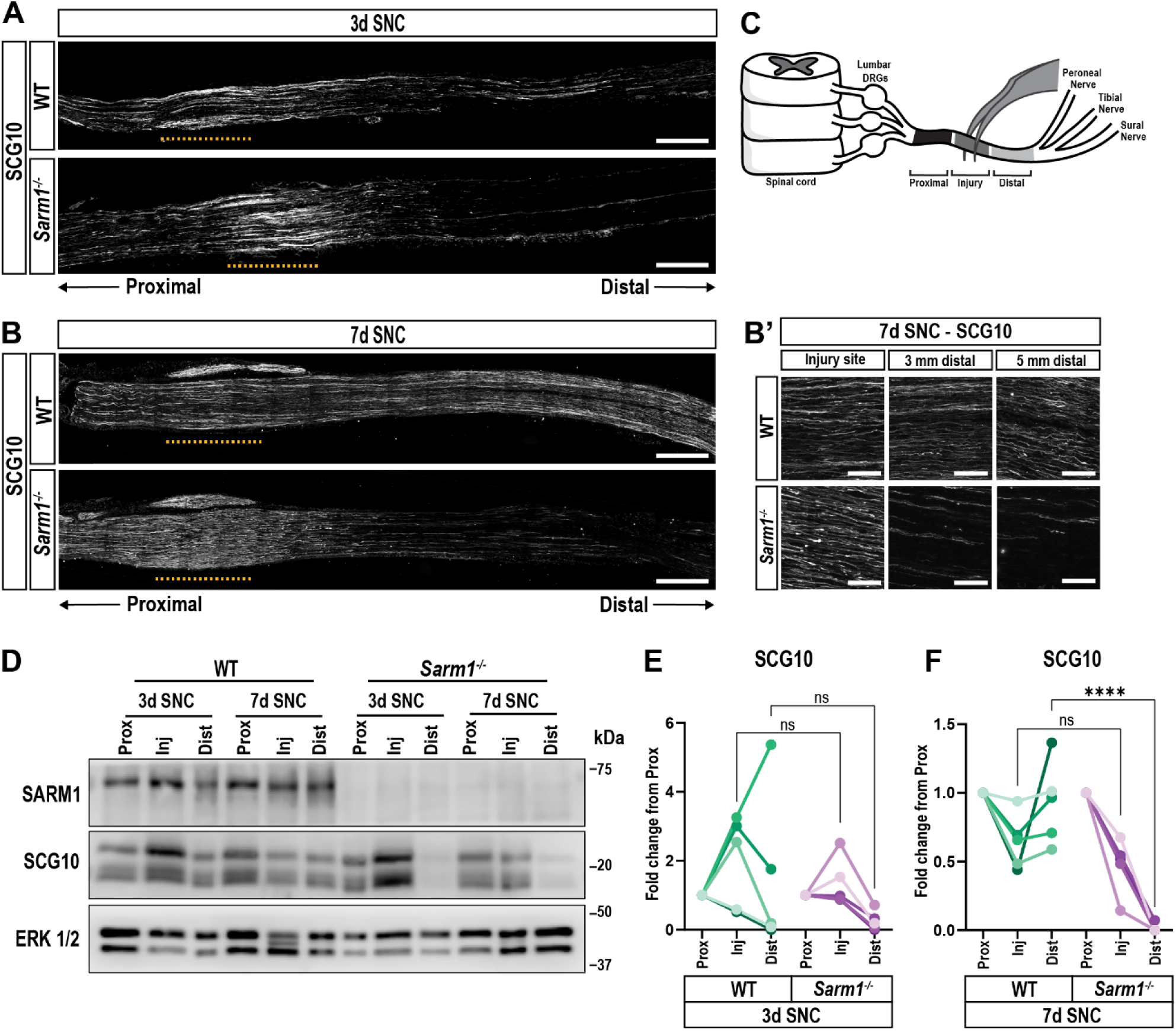
SARM1 is required for the timely regeneration of sensory axons. (**A**) Longitudinal sections of 3 days post-SNC WT and *Sarm1-/-* sciatic nerves stained with anti-SGC10. Yellow dotted lines mark the injury site. Scale bar, 500 µm. (**B**) Longitudinal sections of 7 days post-SNC WT and *Sarm1-/-* sciatic nerves stained with anti-SGC10. Yellow dotted lines mark the injury site. Scale bar, 500 µm. (**B’**) Higher magnification images of SCG10 labeled 7days post-SNC nerves, at the injury site, 3 mm, and 7 mm distal to the injury site. Scale bar, 100 µm. (**C**) Schematic of lumbar spinal cord, DRGs, and sciatic nerve. Brackets depict nerve segments microdissected for Western blotting. (**D**) Immunoblots of nerve segments showing SARM1 and SCG10 in the proximal (Prox), injury site (Inj), and distal (Dist) segments of WT and *Sarm1-/-* mice at 3 and 7 days post-SNC (n = 5 mice per genotype per time point). Anti-ERK1/2 was used as a loading control. (**E** and **F**) Quantification of immunoblots shown in (**D**); n = 5 mice per genotype and time point. Data normalized to ERK1/2 and shown as fold change from proximal nerve for each biological replicate (one-way ANOVA; ****p≤0.0001; ns, not significant).

### *Sarm1* is not required for axotomy-induced activation of DRG neuron intrinsic growth programs

To examine whether delayed regeneration of sensory axons in *Sarm1-/-* mice is due to impaired induction or maintenance of regeneration-associated genes (RAGs), we harvested sciatic DRGs from sham-operated WT and *Sarm1-/-* mice and following injury at 1, 3, and 7 dpc. DRGs were analyzed by bulk RNA-sequencing (**Figure 4A**). In WT DRGs, *Sarm1* expression is not upregulated by nerve injury (**Figure 4B**), and as expected, *Sarm1* is very low in mutant DRGs (**Figure 4B**). Next, we carried out weighted gene co-expression network analysis (WGCNA) (Geschwind and Konopka 2009; Zhang and Horvath 2005). Similar to our previous work (Chandran et al. 2016; Kalinski et al. 2020), we found induction of a pink module, enriched for RAGs (**Figure 4C)**, and a turquoise module, enriched for immune genes (**Figure S4A**).

**Figure 4:**
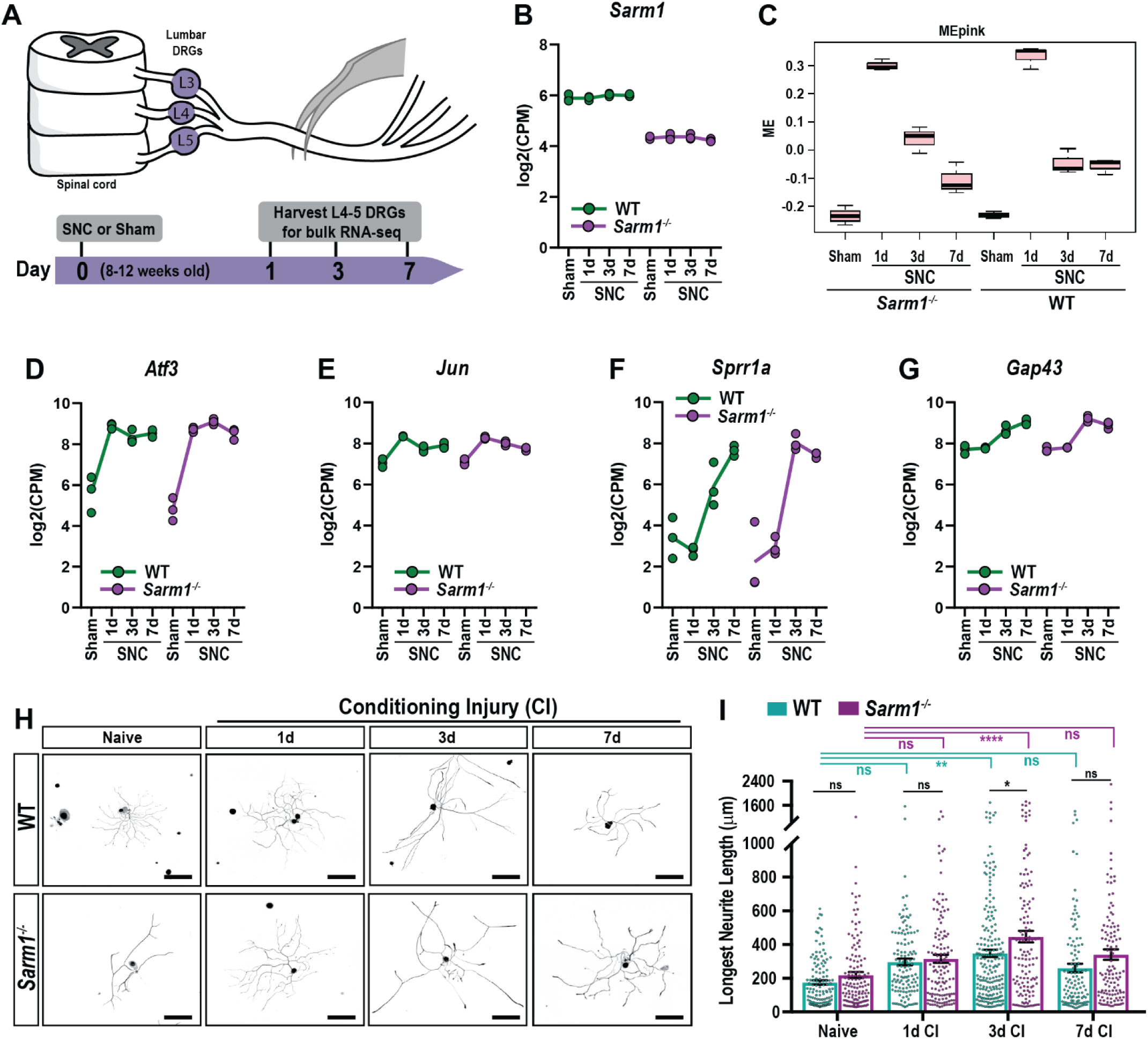
SARM1 is not required for the induction of neuron intrinsic regeneration-associated genes and neurite outgrowth. (**A**) Representation timeline of bulk RNA-seq experiment of sciatic DRGs from sham-operated, 1, 3, and 7 days post-SNC (n = 3 mice per time point and genotype). (**B**) Expression of *Sarm1* in WT and *Sarm1-/-* DRGs, assessed by bulk RNA-seq. CPM, counts per million. (**C**) Weighted gene co-expression network analysis (WGCNA) of WT and *Sarm1-/-* DRGs at different post-injury time points. Module Eigen (ME) scores for the Pink module, enriched for regeneration-associated genes. (**D-G**) Longitudinal expression of the regeneration-associated gene products *Atf3*, *Jun*, *Sprr1a*, and *Gap43* in axotomized DRGs. Solid lines represent the mean of 3 biological replicates. CPM, counts per million. (**H**) Primary DRG neurons prepared from WT and *Sarm1-/-* mice; without or with a 1, 3, and 7-day conditioning injury (CI). Neurons were stained with anti-NF-H (inverted, black). Single neurons were cropped out of a tiled image. Brightness and contrast were adjusted for clarity, using the same parameters for all images. Scale bars, 200 µm. (**I**) Quantification of the longest neurite from cultures shown in (**H**). Each data point equals one neuron (WT and *Sarm1-/-* naïve, n = 247; WT 1d CI, n = 206; *Sarm1-/-* 1d CI, n = 252; WT 3d CI, n =434; *Sarm1-/-* 3d CI, n = 247; WT 7d CI, n = 333; *Sarm1-/-* 7d CI, n = 230). N = 3 biological replicates per group; one-way ANOVA, *p≤ 0.05; **p≤0.01; ****p≤0.0001; ns, not significant.

Baseline module eigen (ME) gene expression for RAGs in the pink module was very similar between WT and *Sarm1-/-* DRGs (**Figure 4C**). At 1 dpc, the pink gene module was upregulated comparably in axotomized WT and *Sarm1-/-* DRGs. At 3 dpc, but not at 7 dpc, the pink module is elevated in *Sarm1-/-* versus WT DRGs (**Figure 4C**). Longitudinal expression analysis of individual RAGs showed comparable induction of *Atf3, Jun, Sprr1a,* and *Gap43,* independently of genotype (**Figure 4D-G**). Injury-regulated gene ontology (GO) terms in WT and *Sarm1-/-* DRGs at 1 dpc include *transsynaptic signaling, cation channel complex, and neuron development*. GO terms at 3 dpc include *biological adhesion, reactome extracellular matrix organization, and epithelial mesenchymal transition*. GO terms at 7 dpc include *reactome ECM proteoglycans, collagen chain trimerization,* and *cell adhesion mediated by integrins*.

The baseline gene expression for the turquoise module is very similar between naïve WT and *Sarm1-/-* mice. However, upon axotomy in WT mice, ME gene expression gradually increases during the first week (**Figure S4A**). In axotomized *Sarm1-/-* mice, there is a massive increase in ME gene expression between 1 and 3 dpc (**Figure S4A**), indicating a more rapid induction of immune genes in axotomized *Sarm1-/-* DRGs. Expression analysis of individual gene products in WT and *Sarm1-/-* DRGs revealed a more rapid increase in *Aif1* (encoding Iba1) and *Itgam* (encoding CD11b) in *Sarm1*-/- compared to WT at 3 dpc (**Figure S4B, C**). Expression levels of *Ccr2* modestly increase and there is no induction of the transcription factors *Stat3*, and *Creb1*, independently of genotype (**Figure S4D-F**). Top upregulated GO terms in the turquoise module include *immune effector process, defense response, innate immune response, and myeloid leukocyte activation*. Collectively, these findings indicate that axotomy triggered upregulation of RAGs and immune genes in sciatic DRGs does occur in *Sarm1-/-* mice, however SARM1 appears to influence the kinetics and magnitude by which these gene modules become activated.

### *Sarm1* is not required for conditioning lesion enhanced axon outgrowth of primary DRG neurons

Next, we assayed axon outgrowth of WT and *Sarm1-/-* DRG neurons *in vitro*. Sciatic DRG neuron cultures were prepared from naïve mice and mice receiving an SNC [conditioning injury (CI)] at 1, 3, and 7 days prior to harvesting of axotomized DRGs. Cultures prepared from naïve DRGs showed comparable axonal length after 20h *in vitro* **(Figure 4H)**. Axon outgrowth was significantly enhanced 3 days following CI in both genotypes, however *Sarm1-/-* neurons showed more growth compared with parallel processed WT neurons **(Figure 4I)**. Taken together, these data show that *Sarm1* is not necessary for upregulation of RAGs in axotomized DRGs or CI-triggered enhancement of axon outgrowth of DRG neurons *in vitro*. Taken together, these findings indicate that reduced axon regeneration in the injured *Sarm1*-/- PNS is not due to failure of activation of neuron-intrinsic growth programs.

### Loss of *Sarm1* delays induction of the SC repair program *in vivo*, but not *in vitro*

Axonal damage causes SC to activate a repair program, including rapid upregulation of sonic hedgehog (*Shh*), c-Jun, and p75^NTR^/NGFR (p75^NTR^). In sciatic nerves of naïve WT and *Sarm1-/-* mice, *Shh* is not detected (**Figure S5A**) and only few p75^NTR^+ non-myelinating (nm)SC are present (**Figure S5B**). Consistent with previous reports, longitudinal sections of injured WT nerves exhibit a strong increase in *Shh* (**Figure 5A-C**) and p75^NTR^ immunoreactivity (**Figure 5D-G**). At 3dpc in WT nerves, *Shh* is upregulated immediately proximal to the crush site, not detected at the crush site, and upregulated throughout the length of the distal nerve. At 3 dpc in *Sarm1-/-* nerves, *Shh* is upregulated immediately proximal to the crush site, not detected at the crush site, and in the distal nerve upregulated only close to the crush site but missing more distally (**Figure 5A, C**). At 7 dpc in WT nerves, *Shh* shows a similar distribution pattern as at 3 dpc, however expression levels are reduced. Interestingly, in 7dpc *Sarm1-/-* mice, strong induction of *Shh* is observed through the entire length of the distal nerve, indicating that *Shh* induction occurs independently of WD(**Figure 5B, C)**. In 3 dpc and 7 dpc WT nerves, strong anti-p75^NTR^ immunoreactivity is observed at the site of nerve injury and within the distal nerve. This stands in marked contrast to parallel processed *Sarm1-/-* nerves, where a strong increase in p75^NTR^ staining is only observed at the nerve injury site, but not in the distal nerve (**Figure 5D, F**). Higher magnification images of the proximal nerve, injury site, and distal nerve show that upregulation of p75^NTR^ at the injury site is *Sarm1*-independent, while the increase in the distal nerve is *Sarm1*-dependent (**Figure 5E, G).** For independent validation of location specific regulation of p75^NTR^ in *Sarm1-/-* nerves by Western blotting, we collected ∼3 mm nerve segments harboring proximal nerve, the injury site, or distal nerve at 3 and 7 dpc (**Figure 3C**). In WT nerves at 3 and 7 dpc, p75^NTR^ is strongly upregulated at the injury site and in the distal nerve (**Figure 5H-J**). In *Sarm1-/-* nerves at 3 and 7dpc, p75^NTR^ is upregulated only at the injury site, but not in the distal nerve (**Figure 5H-J**). In a similar vein, c-Jun upregulation in the distal nerve of *Sarm1-/-* is reduced at 7 dpc compared to WT distal nerve (**Figure 5K, L**). Strong p75^NTR^ expression is sustained in WT distal nerve at 14 and 21 dpc. However, in *Sarm1-/-* distal nerve, a gradual increase in p75^NTR^ is observed (**Figure S5C-E**). Similarly, c-Jun expression is sustained at 14 and 21 dpc in WT distal nerves, and levels are lower in *Sarm1-/-* nerves (**Figure S5C, F, G).** These data indicate that SC reprogramming is delayed in the *Sarm1-/-* distal nerve.

**Figure 5:**
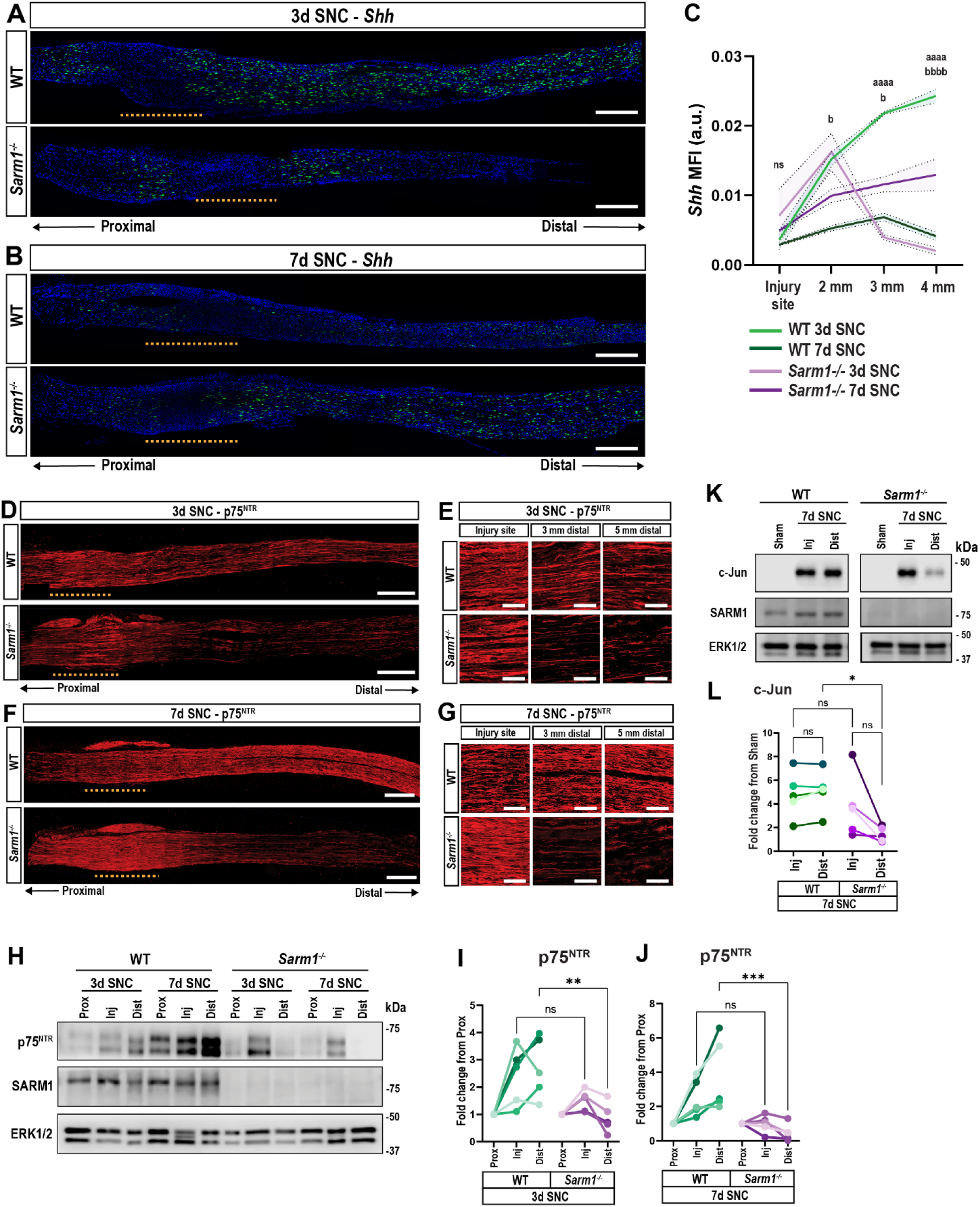
The Schwann cell repair response is delayed in injured *Sarm1-/-* mice. (**A** and **B**) Representative images of longitudinal sciatic nerve sections of WT and *Sarm1-/-* mice at (**A**) 3 days and (**B**) 7 days post-SNC stained for *Shh* transcript (green) and cell nuclei (blue). Yellow dotted lines mark the injury site. Scale bars, 500 µm. (C) Quantification of *Shh* mean fluorescence intensity. Intervals in each graph represent intensity ± SEM. Letter ***a***, comparison between WT and *Sarm1-/-* at 3 days post-SNC; ***b***, comparison between WT and *Sarm1-/-* at 7 days post-SNC. Two-way ANOVA; the number of ***a*** and ***b*** letters is equivalent to the number of asterisks (*p<0.05; ****p<0.0001); a.u., arbitrary units. N = 3 mice per genotype and time point. (D) Representative images of longitudinal sciatic nerve sections of 3 days post-SNC WT and *Sarm1-/-* mice stained with anti-p75^NTR^. Yellow dotted lines mark the injury site. Scale bars, 500 µm; n = 5 mice per genotype. (E) Higher magnification images of p75^NTR^ labeling at the injury site and distal nerve at 3 days post-SNC. Scale bars, 100 µm. (F) Representative images of longitudinal sciatic nerve sections of 7dpc WT and *Sarm1-/-* mice stained with anti-p75^NTR^. Yellow dotted lines mark the injury site. Scale bars, 500 µm; n = 5 mice per genotype. (G) Higher magnification images of p75^NTR^ labeling at the injury site and distal nerve at 7 days post-SNC. Scale bar, 100 µm. (H) Western blots of nerve segments for p75^NTR^ and SARM1. Prox (proximal nerve), Inj (injury site), Dist (distal nerve) of WT and *Sarm1-/-* mice at 3 and 7 days post-SNC (n = 5 mice per genotype; ERK1/2 is shown as loading control). (**I** and **J**) Quantification of Western blots shown in (**H**); n = 5 biological replicates with different color shades. Data normalized to ERK1/2 and shown as fold-change compared to the proximal nerve for each biological replicate (one-way ANOVA; **p≤0.01; ***p≤0.001). (K) Western blots of nerve segments for c-Jun and SARM1. Sham nerves, 7 days post-SNC Inj and Dist segments of WT and *Sarm1-/-* mice are shown (n = 5 mice per genotype; ERK1/2 used as loading control). (L) Quantification of Western blots shown in (**K**); n= 5 biological replicates with different color shades. Data normalized by ERK1/2 and shown as fold-change compared to sham-operated mice. One-way ANOVA; *p<0.05; ns, not significant.

Next, to distinguish between SC intrinsic and extrinsic mechanisms that may underly reduced SC plasticity in injured *Sarm1-/-* mice, we prepared SC cultures from naïve WT and *Sarm1-/-* sciatic nerves and assessed induction of p75^NTR^ and c-Jun *in vitro*. Independently of genotype, p75^NTR^ and c-Jun-labelled SC were abundantly found after 3 days *in vitro* (DIV) (**Figure S5H**). This indicates that *Sarm1* is not required in SC for the generation of rSC *in vitro* and suggests that the microenvironment in the *Sarm1-/-* distal nerve suppresses SC plasticity and activation of the repair program.

### The presence of p75^NTR^+ Schwann cells in the *Sarm1*-/- distal nerve is not sufficient to restore axonal regeneration

To investigate whether delayed SC reprogramming in the *Sarm1-/-* distal nerve is linked to poor regenerative outcomes, we employed a conditioning injury paradigm in which WT and *Sarm1-/-* mice were subjected to two temporally staggered SNCs. A first mid-thigh SNC was followed 10 days later by a second SNC, placed immediately proximal to the first one. After the second SNC, axons were allowed to regenerate for 3 days before nerves were harvested and analyzed (**Figure 6A**). This double (d) crush paradigm, hereafter called dSNC, allows sufficient time for many p75^NTR^ immunoreactive SC to appear in the distal nerve of both, WT and *Sarm1-/-* mice (**Figure 6B**). While dSNC in*Sarm1-/-* mice causes an increase in p75^NTR^ labeling in the distal nerve, levels remained below parallel processed WT mice (**Figure 6B, Figure S5I**). In a similar vein, c-Jun was elevated at the dSNC *Sarm1-/-* injury site, but reduced in the distal nerve, compared to WT dSNC distal nerve (**Figure 6B, S5J).** Western blotting of micro-dissected nerve segments harboring the injury site or distal nerve of dSNC mice (**Figure 6C**) confirmed the more robust increase in p75^NTR^ in WT distal nerve, compared to *Sarm1-/-* distal nerve (**Figure 6C, D**). Similarly, c-Jun expression in the distal nerve of WT mice was significantly higher than in *Sarm1-/-* mice (**Figure 6C, E**). Overall, these data suggest elevated SC plasticity in the *Sarm1-/-* distal nerve following dSNC compared to single SNC, however at a reduced level when compared to parallel processed WT mice.

**Figure 6:**
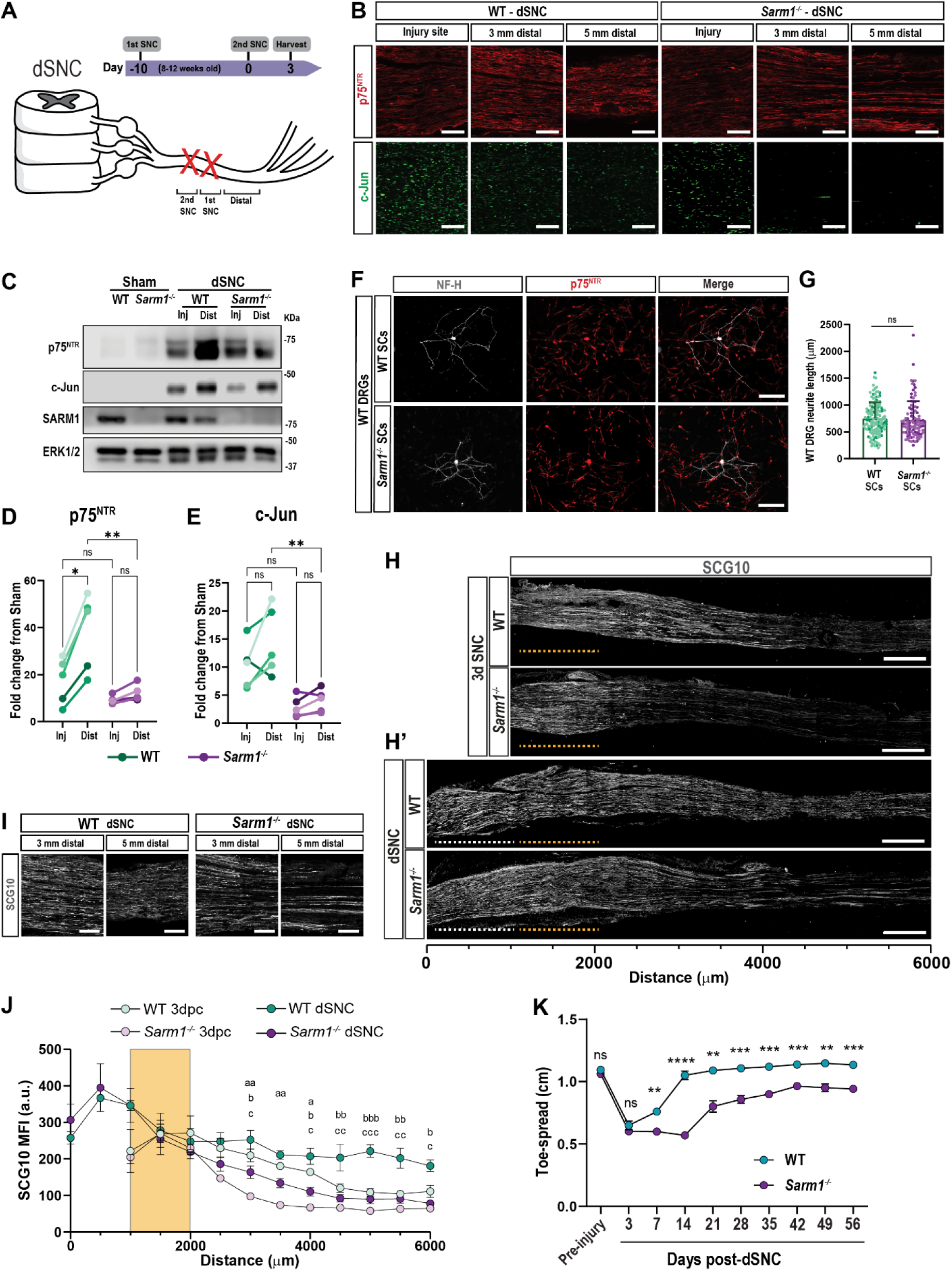
A conditioning lesion is not sufficient to rescue regeneration in *Sarm1-/-* mice. (A) Timeline of the double (d)SNC lesion paradigm. Ten days after the first SNC, a second lesion was placed immediately proximal, and nerves harvested 3 days after the second lesion. (B) Longitudinal sections of dSNC WT and *Sarm1-/-* sciatic nerves showing the injury site and nerve segments 3 mm and 5 mm distal from the injury site, stained for p75^NTR^ (red) and c-Jun (green). Scale bars, 100 µm. (C) Western blots of nerve segments probed for p75^NTR^, c-Jun, and SARM1. WT and *Sarm1-/-* nerves from sham-operated mice, dSNC injury site and distal nerve are shown. ERK1/2 was used as loading control. (**D, E**) Quantification of Western blots shown in (**H**); 5 biological replicates with different color shades. Data was normalized to ERK1/2 and shown as fold change from sham-operated WT or *Sarm1-/-* nerves (one-way ANOVA; *p≤0.05; **p≤0.01); ns, not significant. (F) Anti-NF-H labeled DRG neurons from naïve WT mice co-cultured with SC harvested from WT and *Sarm1-/-* dSNC distal nerves. Scale bars, 250 µm. (G) Quantification of DRG neurite length in cultures shown in (**D**); WT, 188 cells (n = 3 experiments); *Sarm1-/-*, 146 cells (n = 3 experiments). Neurite length ± SEM is shown. Student’s t-test; ns; not significant. (H) Longitudinal sections of 3 days post-SNC (single crush) WT and *Sarm1-/-* nerves stained with anti-SCG10. The yellow dotted lines denote the injury site. Scale bars, 500 µm. (**H’**) Longitudinal sections of dSNC nerves of WT and *Sarm1-/-* mice stained with anti-SCG10. The second crush site is marked with white dotted lines. Scale bars, 500 µm (I) Higher magnification images of SCG10-labeled dSNC WT and *Sarm1-/-* mice at 3 and 5 mm distal to the injury site. Scale bars, 100 µm. (J) Quantification of SCG10 mean fluorescence intensity (MFI) measured at 500 µm intervals from the injury site. Yellow box denotes the first crush site in the dSNC paradigm or the single crush site for the 3 days post-SNC; **a**, WT single crush vs *Sarm1-/-* single crush; **b**, WT single crush vs WT dSNC; **c,** WT dSNC vs *Sarm1-/-* dSNC. MFI ± SEM is shown. Number of ***a***, ***b***, or ***c*** letters is equivalent to the number of asterisks (*p≤0.05; **p≤0.01; ***p≤0.001). N = 3 mice per group. (K) Longitudinal assessment of toe-spread reflex; mean distance between the tips of the first and fifth toes ± SEM is shown. N = 7 mice per genotype and time point. Two-way ANOVA; **p≤0.01; ***p≤0.001; ****p≤0.0001; ns, not significant

As an independent approach to assess SC plasticity in *Sarm1-/-* mice subjected to dSNC, distal nerves were harvested 3 days following the second crush and used for primary SC cultures. Purity of cultured SC was assessed by anti-Sox10 staining. After 1 DIV of both, WT and *Sarm1-/-* cultures, ∼90% of cultured cells were Sox10+, and by 3 DIV ∼80% of cells were Sox10+ (**Figure S6A, B**). Both WT and *Sarm1-/-* SC were able to maintain the rSC phenotype *in vitro*, as evidenced by their elongated morphology and immunoreactivity for p75^NTR^ and c-Jun (**Figure S6C-E**). Next, to assess whether rSC support axon outgrowth, we established co-cultures with WT DRG neurons. Quantification of axon outgrowth revealed comparable lengths between the two genotypes (**Figure 6F, G**). Together, these data suggest that rSC prepared from *Sarm1-/-* and WT dSNC nerve are equally capable of supporting axon outgrowth *in vitro*.

Next, we analyzed axon regeneration following dSNC *in vivo*. In WT mice, dSNC increased sensory axon regeneration when compared to single SNC (**Figure 6H, J**). In *Sarm1-/-* mice however, dSNC was not sufficient to restore regeneration to the level observed in parallel processed dSNC WT mice. When comparing the two genotypes, significantly fewer SCG10+ axons are present in the distal nerve of *Sarm1-/-* mice (**Figure 6H’-J**). This indicates that an increase in p75^NTR^+ SCs in the *Sarm1-/-* distal nerve is not sufficient to rescue sensory axon growth. Commensurate with reduced axon regeneration in *Sarm1-/-* dSNC mice, behavioral studies revealed a significant delay in the recovery of the toe-spread reflex. In WT mice, the reflex is fully restored after 21 days, however in *Sarm1-/-* mice, only partial restoration was observed at 56 days, the latest time point analyzed (**Figure 6K**, **Figure S6F**).

### Loss of *Sarm1* results in a blunted immune response in the distal nerve

Upon SNC, there is a massive infiltration of blood-borne immune cells in the injured nerve (Kalinski et al. 2020). Accumulation of immune cells at the nerve injury site occurs independently of *Sarm1*, while accumulation of macrophages in the distal nerve of injured *Sarm1-/-* mice is significantly reduced (Zhao et al. 2022). To compare the immune milieu in the distal nerve of WT and *Sarm1-/-* mice subjected to dSNC, longitudinal nerve sections were stained for monocyte/macrophage (Mo/Mac) with anti-F4/80 (**Figure 7A, B**), and anti-CD68 (**Figure 7C, D**). Whereas Mo/Mac infiltration to the injury sites was comparable between *Sarm1-/-* and WT, there was a reduction in the distal nerve of dSNC *Sarm1-/-* mice (**Figure 7A-D**). Western blot analysis of dSNC micro-dissected injury and distal nerves further confirmed reduced inflammation in mutants (**Figure 7E, F**). For quantification of Mo/Mac at the injury site and within the distal nerve, we used flow cytometry. While the injury sites of dSNC WT and *Sarm1-/-* mice showed comparable numbers of myeloid cells and lymphocytes, the distal nerve in *Sarm1-/-* mice is significantly less inflamed (**Figure 7G-K**).

**Figure 7:**
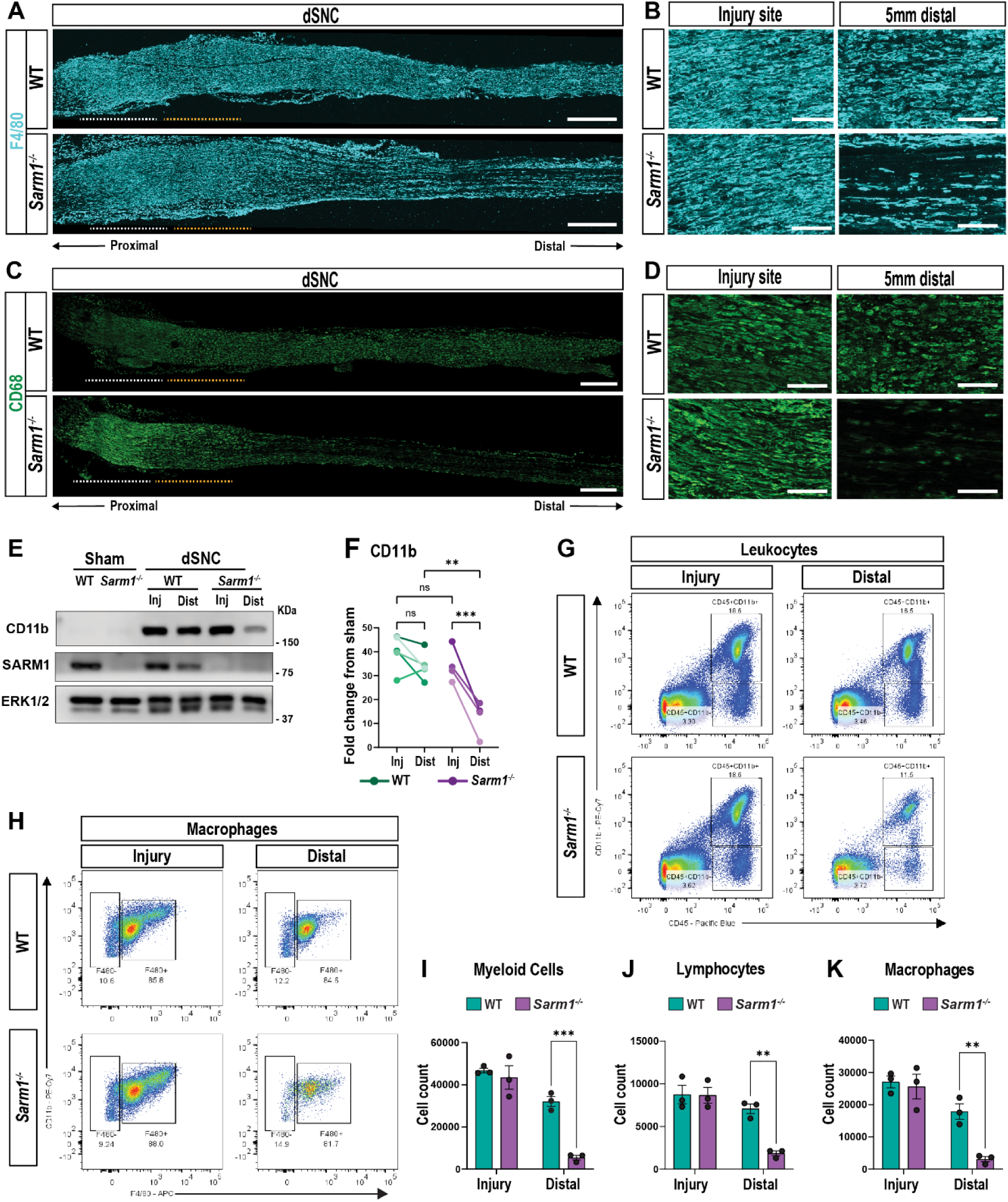
A conditioning lesion is not sufficient to rescue nerve inflammation in *Sarm1-/-* mice. (**A** and **B**) Representative images of F4/80 immunostaining of WT and *Sarm1-/-* nerves subjected to dSNC (n = 3 mice per genotype). First crush site is marked with yellow dotted line; second crush site with white line. Scale bars (**A**), 500 µm; (**B**),100 µm. (**C** and **D**) Representative images of CD68 immunostaining of WT and Sarm1-/- nerves subjected to dSNC (n = 3 mice per genotype). First crush site is marked with yellow dotted line; second crush site with white line. Scale bar (**C**), 500 µm; (**D**),100 µm. (**E**) Western blots of nerve segments probed for CD11b and SARM1. Sham nerves, injury site (Inj) and distal nerve (Dist) of dSNC WT and *Sarm1-/-* nerves are shown. ERK1/2 was used as loading control. (**F**) Quantification of Western blots shown in (**E**); 5 biological replicates with different color shades. Data was normalized to ERK1/2 and shown as fold change from sham-operated WT or *Sarm1-/-* nerves (one-way ANOVA; **p≤0.01; ***p≤0.001; ns; not significant). (**G** and **H**) Flow cytometry dotplots for leucocytes (**G**) and macrophages (**H**) of dSNC WT and *Sarm1-/-* sciatic nerves. Nerves were microdissected into injury site and distal nerves. (**I-K**) Quantification of Myeloid cells (CD11b^hi^ + CD45^hi^), Lymphocytes (CD11b^lo^ + CD45^hi^), and Macrophages (CD11b^hi^ + F4/80^hi^) in dSNC nerves separated into injury site and distal nerve of WT and *Sarm1-/-* mice. N = 3 samples per genotype, with 5 mice per genotype per replica. Flow data are represented as mean ± SEM; two-way ANOVA; **p≤0.01; ***p<0.001.

Since SC and macrophages are important for nerve debridement, we analyzed the integrity of the myelin sheaths in dSNC mice. For lineage tracing and visualization of myelinating SCs (mSCs) we generated *Sarm1+/+; ROSA26^tdT/+^;MPZ^CreER/+^*and *Sarm1-/-;ROSA26^tdT/+^;MPZ^CreER/+^* mice. To induce tdTomato (tdT) expression in mSCs, mice were subjected to tamoxifen (TMX) injection at postnatal day 28 (**Figure S6G**). This approach allows selective labeling of mSCs and does not label other neural crest-derived cells. Tamoxifen-inducible tdT expression in SC was verified by anti-Sox10 staining of naïve sciatic nerves (**Figure S6H**). Upon dSNC, SCs in the distal nerve of WT mice show an elongated morphology consistent with regeneration tracks, whereas most SCs from *Sarm1-/-* resemble mature mSC morphology (**Figure S6I**). **Figure S6J** shows the distal nerves of three biological replicates per genotype; *Sarm1-/-* SCs are consistently less spindled than WT SCs, suggesting that *Sarm1-*dependent Wallerian degeneration is necessary for the formation of Bands of Büngner.

We have previously shown that myelin clearance is compromised in *Sarm1-/-* distal nerve at 7 dpc (Zhao et al. 2022). Upon dSNC, myelin is efficiently cleared from the injury site in both WT and *Sarm1-/-* mice. In the distal nerve, myelin remains present in *Sarm1-/-* mice, but has largely disappeared from parallel processed WT nerves (**Figure 8A, B**). To validate delayed myelin clearance, we performed Western blotting for myelin basic protein (MBP) and myelin protein zero (P0) using nerve segments harvested from the injury site and the distal nerve, respectively (**Figure 8C-E**). In the distal nerve of dSNC WT mice, myelin proteins are strongly reduced compared to naïve nerve, however in dSNC *Sarm1-/-* mice, myelin proteins are abundantly detected (**Figure 8D, E).**

**Figure 8:**
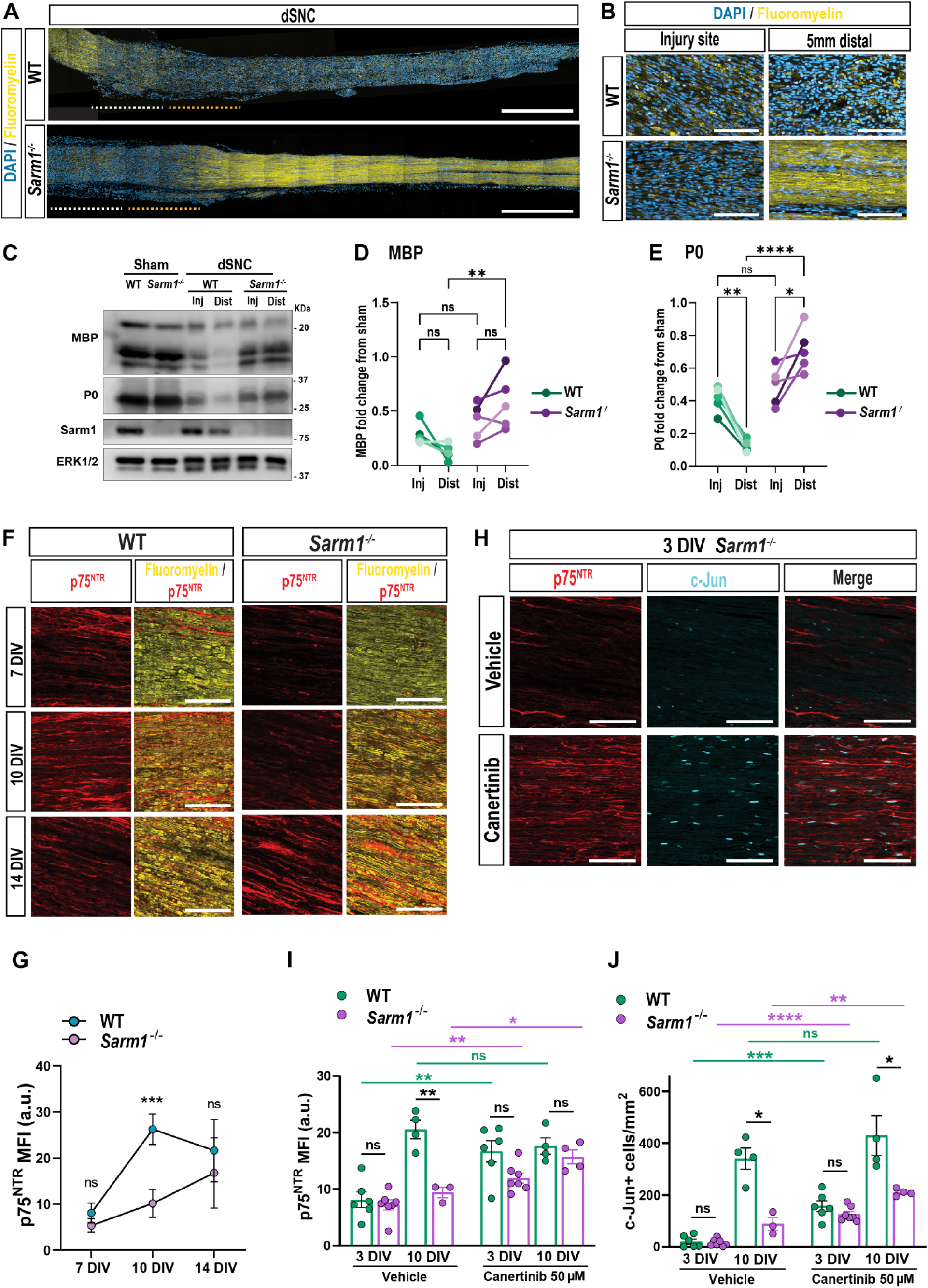
Inhibition of ErbB signaling in *Sarm1-/-* nerves results in rapid activation of the Schwann cell repair response. (**A** and **B**) Longitudinal nerve sections of dSNC WT and *Sarm1-/-* mice stained with fluoromyelin (yellow) and cell nuclei (DAPI, blue). First crush site is marked with yellow dotted line; second crush site with white dotted line. N = 5 mice per genotype. Scale bar, 1000 µm. (**C**) Western blots of nerve segments probed for myelin basic protein (MBP), myelin protein zero (P0), and SARM1. The injury site and distal nerve segments of WT and *Sarm1-/-* mice are shown, along with nerves from sham-operated mice for comparison. ERK1/2 was used as loading control. (**D** and **E**) Quantification of Western blots shown in (**C**); n = 5 biological replicates with different color shades. Data was normalized to ERK1/2 and shown as fold change compared to sham nerves (one-way ANOVA; *p≤0.05; **p≤0.01; ****p≤0.0001; ns, not significant). (**F**) *Ex vivo* cultures of WT and *Sarm1-/-* sciatic nerve trunks, sectioned after 7, 10, and 14 days *in vitro* (DIV), and stained with fluoromyelin and anti-p75^NTR^. Scale bars, 100 µm. (**G**) Quantification of labeled sections shown in (**F**). p75^NTR^ mean fluorescence intensity (MFI); a.u., arbitrary units ± SEM. (n = 4 mice per genotype; Student’s t-test, ***p≤0.001; ns, not significant). (**H**) *Ex vivo*, *Sarm1-/-* nerves were cultured for 3 DIV in the presence of vehicle or Canertinib (50 µM). Nerves were stained for p75^NTR^ and c-Jun. Scale bar, 100 µm. (**I** and **J**) Quantification of staining from *ex vivo* nerves treated with vehicle or Canertinib for 3 and 10 DIV. p75^NTR^ MFI (**I**) and number of c-Jun+ nuclei per mm^2^ ± SEM (**J**). a.u., arbitrary units; n = 3-7 mice per genotype and time point; Two-way ANOVA for comparisons across genotypes and Student’s t-tests for comparisons within the same genotype; *p≤0.05; **p≤0.01; ****p≤0.0001; ns, not significant.

To ask whether delayed SC plasticity and myelin clearance in *Sarm1-/-* mice is due to absence of blood-derived immune cells, we prepared *ex vivo* nerve cultures from naïve WT and *Sarm1-/-* mice and assessed myelin clearance after 7, 10, and 14 DIV (**Figure 8F**). Independent of genotype, myelin clearance was protracted, as assessed by fluoromyelin staining. Moreover, compared to WT, the appearance of p75^NTR^+ rSCs is delayed in *Sarm1-/-* nerves *ex vivo* (**Figure 8F, G**). This shows that the reprograming of SC into rSC does not require nerve infiltration of blood-derived immune cells, and thus, delayed SC reprogramming in the *Sarm1-/-* distal nerve is likely not due to reduced accumulation of macrophages. The studies also underscore the importance of macrophages for rapid myelin clearance.

### Inhibition of ErbB induces the SC repair program in *Sarm1-/-* nerves

Neuregulin-1 type III (NRG1) is located on the axonal surface where it interacts with the epithelial growth factor receptor (EGFR) family members ErbB2 and ErbB3 produced by SC (Zhao et al. 2022). NRG1 regulates SC differentiation and is important for induction and maintenance of the myelinating state (Quintes et al. 2010; Daboussi et al. 2023). Because the SC repair program is not effectively activated in *Sarm1-/-* nerves *ex vivo*, this provided an opportunity for pharmacological examination of signaling pathways that suppress induction of the repair program. To test whether prolonged NRG1-ErbB2/3 signaling in *Sarm1-/-* nerves underlies the protracted induction of c-Jun and p75^NTR^, cultured nerves were treated with vehicle or Canertinib, a pan-ErbB receptor tyrosine kinase inhibitor. After 3 and 10 DIV, nerves were collected, sectioned, and stained with anti-p75^NTR^ and anti-c-Jun. While p75^NTR^ and c-Jun were low in vehicle-treated nerves at 3 DIV, a significant upregulation was observed in Canertinib-treated cultures (**Figure 8H-J**). At 10 DIV, p75^NTR^ and c-Jun were significantly elevated in vehicle treated WT nerves, compared to *Sarm1-/-* nerves. However, Canertinib treatment of *Sarm1-/-* nerves resulted in a significant increase in p75^NTR^ and c-Jun. While p75^NTR^ levels in Canertinib treated *Sarm1-/-* nerves are comparable to WT nerves, c-Jun did not reach WT levels (**Figure 8I, J**). Our findings suggest that inhibition of ErbB kinase in *Sarm1-/-* nerves *ex vivo* mimics activation of the SC repair program.

### The *Sarm1* nerve microenvironment is not conducive for axon regeneration

To directly demonstrate that the *Sarm1-/-* nerve microenvironment fails to support robust regeneration of severed WT axons, we carried out nerve grafting experiments. In a first set of experiments, we grafted acutely isolated WT sciatic nerve trunks into WT recipients (WT → WT) by suturing the epineurium of the graft to the epineurium of the proximal nerve stump of the host (**Figure 9A)**. We then determined the time it takes to observe robust growth of SCG10+ axons into WT nerve grafts and found 14 days as a suitable time point (data not shown). Next, we grafted acutely isolated *Sarm1-/-* nerves into WT hosts (*Sarm1-/-* ◊ WT) and assessed axon regeneration into the graft. After 14d, myelin ovoids have formed in WT and *Sarm1-/-* grafts, however, clearance of myelin is delayed in *Sarm1-/-* grafts (**Figure 9B**), likely due to the reduced accumulation of macrophages (**Figure S9A, B**). After 14 days, significantly fewer SCG10+ axons regenerated into *Sarm1-/-* grafts compared to WT grafts (**Figure 9C-E**). Moreover, p75^NTR^ staining was reduced in *Sarm1-/-* grafts (**Figure 9C, D, F**). Together these experiments show that WT leukocytes fail to enter the *Sarm1-/-* graft and that WT DRG neurons fail to regenerate axons into the *Sarm1-/-* graft. We conclude that SARM1 function in the distal nerve is necessary for the timely regeneration of sensory axons.

**Figure 9:**
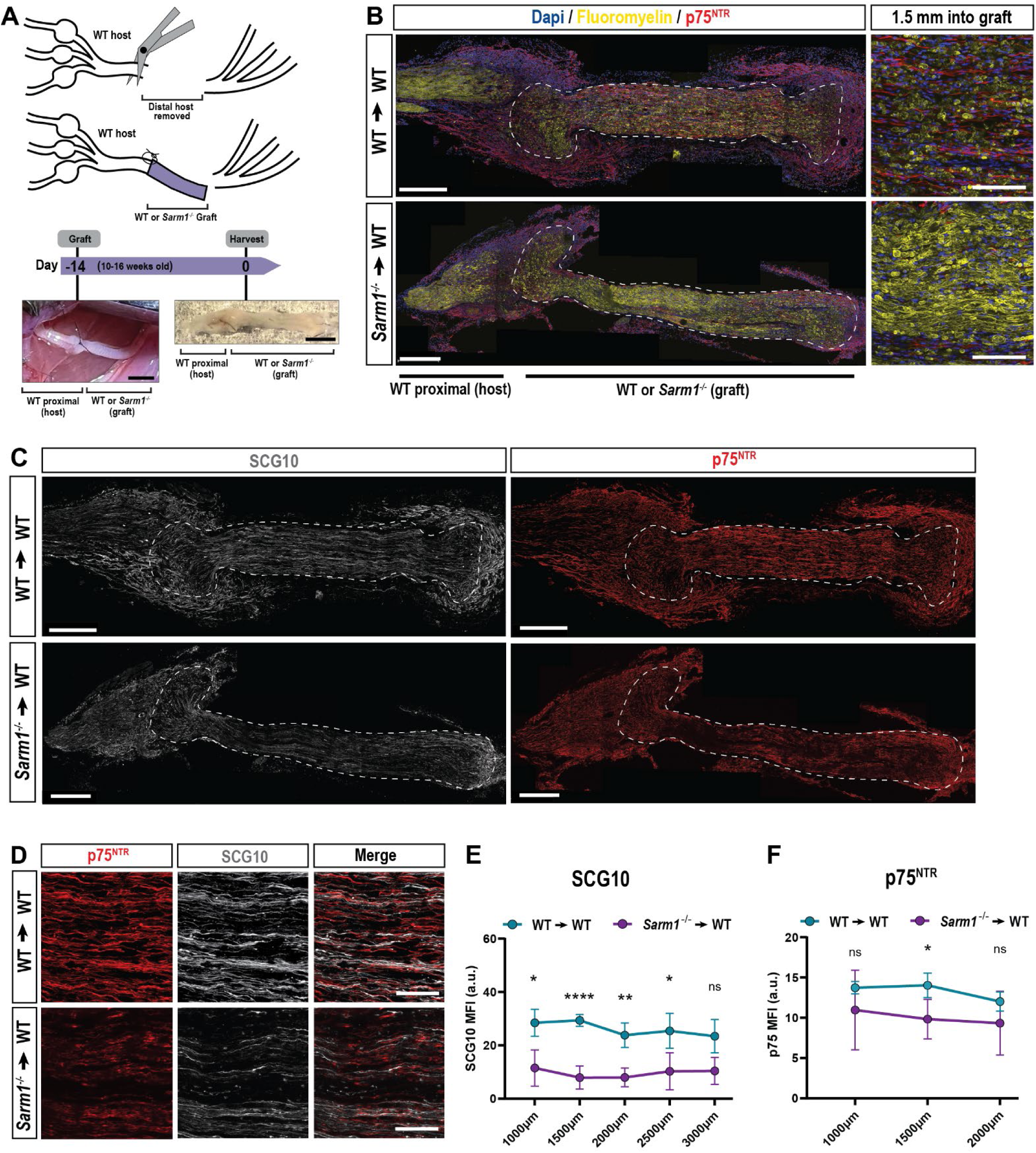
*Sarm1-/-* nerve microenvironment is not conducive for nerve regeneration. (**A**) Schematic of the sciatic nerve grafting experiment. A WT or *Sarm1-/-* nerve segment was grafted into a WT host. Coaptation of the graft was done only at the proximal end. Intraoperative and harvested graft images are shown. Scale bars, 1 mm. (**B**) Representative longitudinal nerve sections showing WT and a *Sarm1-/-* grafts (dotted lines) after 14 days, stained with fluoromyelin (yellow), cell nuclei (DAPI, blue), and anti-p75^NTR^. High magnification images on the right show difference in myelin integrity between WT and *Sarm1-/-* grafts. Scale bars for low magnification, 500 µm; scale bars for high magnification, 100 µm. (**C**) Nerve sections of WT and *Sarm1-/-* grafts stained with anti-SCG10 and anti-p75^NTR^. (**D**) High magnification of WT and *Sarm1-/-* grafts stained with anti-p75^NTR^ and anti-SCG10. Scale bars, 100 µm. (**E** and **F**) Quantification of p75^NTR^ and SCG10 mean fluorescence intensity ± SEM at 500 µm intervals within the grafts (n = 4-6 biological replicates per genotype). Student’s t-tests at each individual distance; *p≤0.05; **p≤0.01; ****p≤0.0001; ns, not significant.

## Discussion

Our studies demonstrate that *Sarm1*-dependent WD in the PNS is necessary for the timely regeneration of sensory and motor axons. Regenerated nerve fibers in *Sarm1-/-* mice exhibit a long-lasting reduction in axon caliber, coinciding with a smaller amplitude of compound action potentials. Defective nerve regeneration results in delayed restoration of the toe-spread reflex and skilled hind foot placement. Mechanistically, *Sarm1* is not necessary for SNC-elicited activation of neuron-intrinsic growth programs in axotomized DRGs or the initiation of regenerative axon growth into the nerve injury site. In the distal nerve, however, *Sarm1* is necessary for the rapid activation of the SC repair program, nerve inflammation, clearing of myelin debris, and axonal growth. Grafting *Sarm1-/-* nerve into WT recipient mice provides independent evidence that blocking WD results in an adverse nerve microenvironment, inhibitory for WT axon regeneration. Based on these observations, we propose that SARM1 function in distal axons, but not in neuronal cell soma or proximal axons, is necessary for timely PNS repair.

Nerve grafting studies in rodents, using pre-degenerated nerve sutured to an acutely injured sciatic nerve, found superior axon regeneration compared to grafting of freshly harvested nerve (Danielsen et al. 1994). However, the underlying cellular and molecular mechanisms remained incompletely understood. Studies with *Wld^S^* mice and zebrafish neurons overexpressing *Wld^S^* indicate that WD-resistant axons inhibit regeneration (Martin et al. 2010). Moreover, in *Wld^S^* mice, regeneration of sensory axons is significantly more impeded than motor axons, arguing against simple physical obstruction (Brown, Lunn, and Perry 1992; Lunn et al. 1989). In a similar vein, axon regeneration in *Sarm1-/-* mice is delayed, for both, sensory and motor axons. Delayed axon regeneration in *Sarm1-/-* mice is not due to failure of injury-induced activation of neuron intrinsic growth programs, as demonstrated by upregulation of RAGs in axotomized DRGs, and enhanced axon outgrowth of primary DRG neurons subjected to a conditioning lesion. We find that following a 3-day conditioning lesion, axon outgrowth in *Sarm1-/-* cultures is increased compared to parallel processed WT cultures, suggesting that neuronal *Sarm1* may inhibit axon outgrowth. Commensuarate with this, studies in invertebrates found that *Sarm1/Tir-1* functions cell-autonomously to inhibit axon regeneration (Czech et al. 2023). Noteworthy, our findings are at variance with primary DRGs neurons from *Wld^S^* mice, where conditioning lesion enhanced axon outgrowth was significantly reduced (Niemi et al. 2013), indicating there may be important differences between axotomized *Wld^S^* and *Sarm1-/-* sensory neurons.

We previously showed that in WT and *Sarm1-/-* mice subjected to SNC, the lesion site is rapidly cleared of myelin debris and that blood-borne leukocytes accumulate at comparable numbers (Zhao et al. 2022). In PNS injured *Sarm1-/-* mice, sensory axons are capable of rapid axon extension into the lesion site, similar to WT mice, providing additional evidence that activation of neuron intrinsic growth programs remains intact. In marked contrast, during the first week post-SNC, sensory axon growth into distal nerve is only observed in WT but not *Sarm1-/-* mice. Moreover, a conditioning lesion in *Sarm1-/-* mice is not sufficient to rescue the delayed axon regeneration in the distal nerve *in vivo*. The poor regenerative growth of sensory axons in the *Sarm1-/-* distal nerve coincides with a delayed and more gradual appearance of rSC versus the acute appearance of rSC in the injured WT PNS. We speculate that the slow disintegration of severed axons in the *Sarm1-/-* distal nerve results in a gradual drop of axonal signals that normally keep SC in a differentiated state. And as a result, induction of the SC repair program is not only delayed but also blunted. In support of this idea, the actual process of axon destruction, when it does occur in *Wld^S^*, is different from WD observed in WT mice (Beirowski et al. 2005). Studies with *Wld^S^*mice revealed that axon degeneration is not only delayed, but that disintegration occurs through a fundamentally different mechanism. WD resistant axons in *Wld^S^* mice undergo a slow anterograde decay, presumably because vital axonal components are gradually depleted. Thus, if WD is blocked, axons do not simply follow the degeneration pathway of WT axons at a slower rate (Beirowski et al. 2005). We provide evidence that prolonged signaling of axonal NRG1 to ErbB on the surface of mature SC in *Sarm1-/-* nerves blocks the induction of the SC repair response. Because ErbB inhibition results in rapid upregulation of c-Jun in *Sarm1-/-* nerves, prior to axon degeneration, this suggests that *Sarm1-/-* SC are capable of rapid activation of the repair program.

Because of the central role of denervated SC in shaping the nerve microenvironment and axon regeneration, alterations in SC plasticity in *Sarm1-/-* mice have far-reaching consequences. NRG1 drives SC differentiation and myelin gene expression; prolonged ErbB signaling may contribute to the failure to downregulate myelin gene products, fragmentation of myelin sheaths, and myelin autophagy (Belgrad, De Pace, and Fields 2020; Jessen and Mirsky 2016). A compounding effect on myelin clearance is the protracted accumulation of macrophages in the *Sarm1*-/- distal nerve. Macrophage phagocytosis plays a vital role in myelin removal, especially during the later phase of myelin clearing. Because some myelin-associated proteins strongly inhibit axon outgrowth (McKerracher et al. 1994; Schäfer et al. 1996), slow myelin clearance likely contributes to delayed regeneration (Yuan et al. 2022).

There is ample evidence that extrinsic factors, in the local nerve microenvironment, can profoundly influence the success of axon regeneration (Rigoni and Negro 2020). For example, the regenerative capacity of the injured mammalian PNS strongly declines with age (Nagano 1998; Verdú et al. 2000). Mechanistic studies revealed an age-related decline in WD, but not in injury-induced activation of regeneration-associated transcriptional programs in axotomized neurons (Painter et al. 2014). The same study found diminished SC repair responses in the distal nerve. Whether the delay in SC reprogramming reflects delayed WD in aged mice, or a diminished capacity of denervated SC to reprogram, remains unknown. Similar to aged WT mice, injured young adult *Sarm1-/-* mice show greatly delayed WD, and this delay blunts the SC repair response. Thus, our studies show that a delay in WD is sufficient to impair reprogramming of young SC. This also shows that an unfavorable nerve microenvironment can suppress regenerative axonal growth even if neuron intrinsic growth programs are properly activated. Additional studies are needed to determine whether the inhibition of ErbB signaling in injured *Sarm1-/-* mice or aged WT mice allows a more rapid activation of the SC repair program and improved axon regeneration.

The similarities in the injury responses between aged WT mice and young *Sarm1-/-* mice begin to shed light on the physiological significance of why axons possess an evolutionarily conserved and highly effective ‘self-destruct’ machinery. Rapid *Sarm1*-dependent WD is crucial for shaping of the distal nerve microenvironment, successful axon regeneration, and the restoration of motor function. While improved axon protection is desirable to shield against nerve damage inflicted by chemotherapy, pollutants, metabolic disorders, or genetic predisposition, it may be counter indicative for axonal regeneration, where blocking the SC repair response is likely detrimental.

## Material and Methods

### Mice and genotyping

All procedures involving mice were approved by the local Institutional Animal Care and Use Committee (IACUC) and performed in accordance with guidelines developed by the National Institutes of Health. Adult (8– 15-week-old) male and female mice on a C57BL/6 background were used throughout the study. Transgenic mice included, *Sarm1/Myd88-5-/-* (Jackson labs stock # 018069), *MPZ-Cre^ERT2^* (Leone et al., 2003), and ROSA26Sortm34.1(CAG-Syp/tdTomato)Hze/J (Jackson labs stock #012570). Mice were housed under a 12 h light/dark cycle with chow and water ad libitum. For genotyping, tail or ear biopsies were collected and genomic DNA extracted by boiling in 100 µl alkaline lysis buffer (25 mM NaOH and 0.2 mM EDTA in ddH_2_O) for 30 min. The pH was neutralized with 100 µl of 40 mM Tris-HCI (pH 5.5). For PCR, 1–5 µl of gDNA was mixed with 0.5 µl of 10 mM dNTP mix (Promega, C1141, Madison, WI), 10 µl of 25 mM MgCl_2_, 5 µl of 5X Green GoTaq Buffer (Promega, M791A), 0.2 µl of GoTaq DNA polymerase (Promega, M3005), 1 µl of each PCR primer stock (100 µM each), and ddH_2_O was added to a total volume of 25 µl. The following PCR primers, purchased from *Integrated DNA Technologies,* were used (**Table 1**). PCR conditions were as follows: Hot start 94°C 3 minutes; DNA denaturing at 94°C 30 seconds; annealing 60°C 1 minute; extension 72°C 1 min, total cycles 34. Final extension for 6 min at 72°C.

**Table 1:**
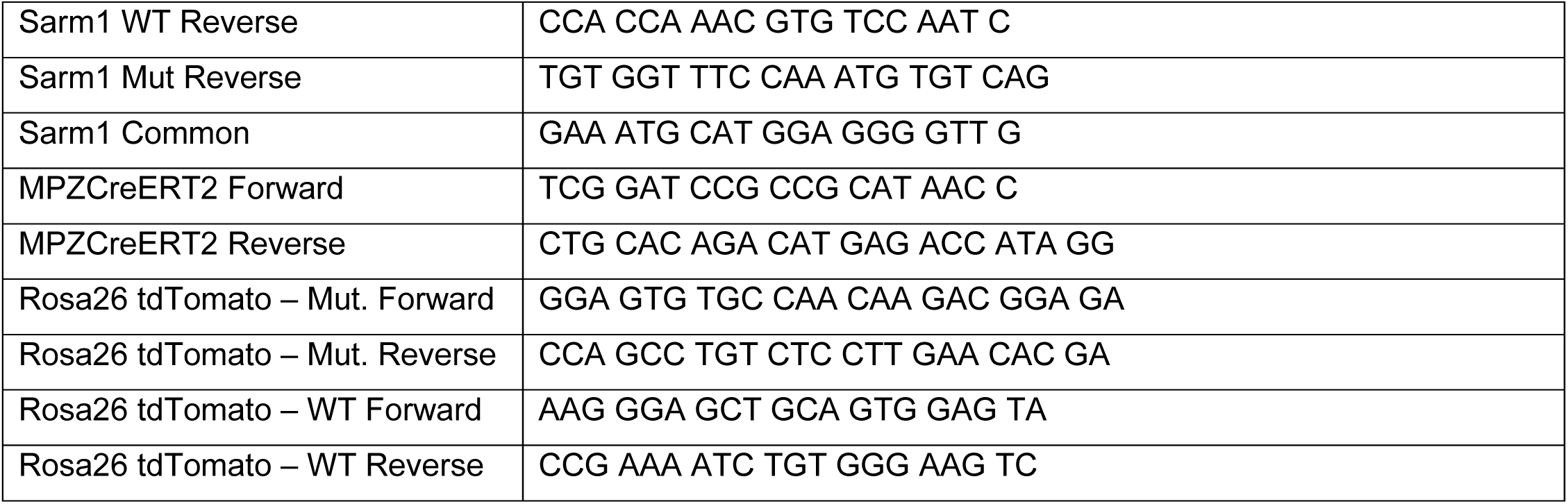
Primer sequences:

### Surgical procedures

Mice were deeply anesthetized with a mixture of ketamine (100 mg/kg) and xylazine (10 mg/kg) or with isoflurane (5% induction, 2-3% maintenance, SomnoSuite Kent Scientific). Buprenorphine (0.1 mg/kg) was given as an analgesic. For sciatic nerve surgery, thighs were shaved and disinfected with 70% ethanol and iodine (PDI Healthcare). A small incision was made on the skin, underlying muscles separated, and the sciatic nerve exposed. For sham operated mice, the nerve was exposed but not touched and the wound closed. For SNC, the nerve was crushed at mid-thigh level for 15 seconds, using a fine forceps (Dumont #55, FST). For SNT, the nerve was cut with neurosurgical microvascular STA-MCA scissors (Codman 80-1803 Scissors, 60° Angled, straight; Aspen Surgical, Caledonia, MI). For nerve grafting experiments, a mid-thigh nerve segment, approximately 7-10 mm in length was acutely isolated from a donor mouse and kept on ice in PBS until ready for grafting. From the recipient mouse, a segment of the sciatic nerve trunk was removed. Coaptation of the graft onto the proximal nerve stump of the recipient was done by suturing the epineurium with a single 10/0 nylon suture (Fine Science Tools, S&T sutures). Wounds were closed using 7mm reflex clips (Cell Point Scientific) and clips removed after 7d. Nerves were harvested 14d after grafting.

Conditioning lesion of sciatic nerve. A first SNC was applied as described above. To perform the second SNC in the double (d)SNC paradigm, mice were anesthetized with isoflurane (5% induction, 2-3% maintenance, SomnoSuite Kent Scientific) 10d after the first SNC and the crush placed immediately proximal to the first one. Wounds were closed with 7mm reflex clips (Cell Point Scientific) and sciatic nerves were harvested 3d after the second SNC.

### Immunofluorescence staining

Animals were deeply anesthetized with a mixture of ketamine/xylazine (25 mg/ml ketamine and 2.5 mg/ml xylazine in PBS) and perfused trans-cardially with ice-cold PBS for 5 min, followed by ice-cold 4% paraformaldehyde in PBS (4%PFA/PBS) for 5 min. Sciatic nerves, DRGs, and spinal cord were dissected and post fixed for 2-4 hours in 4%PFA/PBS. Tissues were transferred to 30% sucrose in PBS incubated at 4°C for at least 24 hours, embedded in OCT (Tissue-Tek, 4583) and frozen on crushed dry ice. Nerves and DRGs were serially sectioned at 12-14 µm thickness at −20°C, using a Leica CM 3050S Cryostat. Sections were mounted onto Superfrost Plus (Fisherbrand, 12-550-15) or Superfrost Plus Gold (Fisherbrand, 15-188-48), microscope slides, air dried overnight, and stored at −20°C.

Sections were warmed to RT, rinsed twice in 1X PBS for 5 minutes each and permeabilized in 0.3% triton-x-100/PBS for 10 minutes. For blocking, sections were incubated in 5% donkey serum (EMD Millipore, S30) in 0.1% Triton-x-100/PBS for 1 hour. For staining the following primary antibodies were used: rabbit α-STMN2/ SCG10 (Novus Biologicals, NBP149461, 1:2000), α-STMN2/SCG10 (Thermo Fisher Scientific, 720178, 1:1000); rat α-F4/80 (Thermo Fisher Scientific ma1-91124, 1:500), CD68, α-Iba1 (Wako Chemicals, 019-19741, 1:1000), goat α-p75^NTR^ (Neuromics GT15057, 1:1000); chicken α-neuofilament H (Aves Lab NFH, 1:1000), and c-Jun (Cell Signaling #9165, 1:200), in blocking buffer overnight at 4°C in a humidified chamber. The next day, sections were rinsed three times in 1X PBS, 5 minutes each and were incubated in secondary antibodies (donkey anti-chicken cy3 (Jackson Immunoresearch, 703-165-155); donkey anti-rat Alexa 488 (Life Technologies, A21208); donkey anti-rabbit cy5 (Jackson Immunoresearch, 711-175-152), donkey anti-goat DyLight 405 (Jackson Immunoresearch, 705-475-003), all at 1:500, in blocking buffer for 1 hour in a humidified chamber. Fluoromyelin stain (Thermo-Fisher, F34651) was conducted after secondary antibodies for 20 min (1:300 diluted in 1x PBS). Sections were mounted in ProLong Gold (Invitrogen, P36930) anti-fade mounting medium and imaged. Sciatic nerve sections were imaged by an experimenter blinded to mouse genotype using a Zeiss Axio Observer. Z1 inverted fluorescence microscope fitted with a Zeiss Axiocam 503 mono camera. The Apotome was enabled for all images at a scan setting of 3 images per z plane. Whole nerve sections were tiled scan using Zen Blue autofocus software in a single plane with a HC PL APO 20x/0.75 objective. High magnification, single tile images were taken on an HC PL APO 63x/1.40 objective. Optical sections were taken through the tissue at 1.0 µm z-step size.

Mean fluorescence intensity of SCG10 in SNC and dSNC was measured using ImageJ or Rstudio at 500 μm intervals from the start of the injury site (identified using F4/80 staining) within a 50 μm wide area set to the thickness of the nerve. SCG10 intensity in the grafts was measured at 1000 µm from the end of the host proximal stump, and at 500 µm intervals as described above. Mean intensity of p75^NTR^ and Iba1 in the grafts and in the in vitro nerve explants was measured using ImageJ at 500 µm intervals within a 300 µm^2^ area.

### RNA in situ hybridization

Mice were perfused for 5 min with ice-cold PBS, followed by RT 10% Neutral Buffered Formalin (NBF, Fisher Chemical, SF-100). Nerves were harvested and postfixed in 10% NBF overnight at RT and transcripts detected with the RNAscope Multiplex Fluorescent Reagent Kit v2 (ACD, 323100). Microscope slides with serially sectioned nerves were rinsed in 1× PBS for 5 min (repeated twice) and air-dried by incubation at 70°C for 10 min in an oven (VWR Scientific, Model 1525 incubator). Next, tissue sections were post-fixed in ice-cold 10% NBF for 15 min and dehydrated by incubation in a graded series of 50%, 70%, and 100% ethanol for 5 min each. Sections were air-dried for 5 min at RT and one drop of hydrogen peroxide solution (ACD Cat# PN 322381) was added to each nerve section on each slide and incubated at RT for 10 min. Sections were then submerged in 99°C RNase-free water for 15 s, followed by incubation in 99°C 1× antigen retrieval solution (ACD Cat# 322000) for 6 min. The slides were washed in MilliQ water twice, followed by a transfer to 100% ethanol for 3 min at RT. The slides were taken out and air-dried. Protease Plus solution (ACD Cat# PN 3223311) was applied to tissue sections followed by incubation at 40°C in an ACD hybridization oven (ACD Cat# 321710) for 10 min followed by MilliQ H2O washes. RNA probes were mixed at appropriate ratios and volumes (typically 50:1 for C1:C2) for complex hybridization. For single RNA probe hybridization, RNA probes were diluted with probe dilutant at 1:50– 1:100 (ACD Cat# 300041). Appropriate probes or the probe mixtures were applied to tissue sections and incubated for 2 hr in the hybridization oven at 40°C. A 1× wash buffer solution was prepared from a 50× stock (ACD Cat# PN 310091) and sections rinsed for 2 min. Amplification probe 1 was applied and slides incubated in a hybridization oven for 30 min 40°C and rinsed twice with 1× wash solution. Next, the A2 and A3 probes were applied. For development, the TSA system (AKOYA, Cy3: NEL744001KT; Cy5: NEL745001KT; Fluorescein: MEL741001KT) was used. Once the color for probe C1 was selected, HRPC1 solution (ACD Cat# 323120), it was applied to the appropriate sections and incubated for 15 min in the hybridization oven at 40°C. The sections were then rinsed in 1× wash solution. Designated TSA color for probe C1, diluted in the TSA dilutant (ACD Cat# 322809) at 1:2000 was applied to the respective sections and incubated for 30 min in the ACD hybridization oven at 40°C. Sections were rinsed in 1× wash solution and then HRP blocker (ACD Cat# 323120) was applied and incubated for 15 min in the ACD hybridization oven at 40°C. This procedure was repeated for probes C2 and C3 as needed using HRPC2 and HRPC3, respectively. Sections were mounted in DAPI Southern Biotech mounting media (Cat# 0100-20), air-dried, and imaged and stored at 4°C in the dark. For quantification of labeled cells in nerve tissue sections, an FoV was defined, 100 µm × 100 µm at the nerve injury site and at multiple distances in increments of 1000µm from the injury site, in the distal nerve. Mean Fluorescent Intensity per FoV was counted. Only labeled cells with a clearly identifiable nucleus were included in the analysis. The Mean Fluorescent Intensity measured per FoV was from n = 3 mice with n = 2 FoV per distance point within each nerve.

### Toluidine blue staining and transmission electron microscopy

The main trunk of the sciatic nerves from naïve WT and *Sarm1-/-* mice was used for combined light microscopy (LM) and ultrastructural analysis and compared to mice 49 days after a mid-thigh sciatic nerve crush injury. Animals were deeply anesthetized with a mixture of ketamine/xylazine (25 mg/ml ketamine and 2.5 mg/ml xylazine in PBS) and perfused trans-cardially at a rate of 1.5mL/10 minutes with ice-cold PBS for 1 min, followed by ice-cold 2% glutaraldehyde, 2% paraformaldehyde in phosphate buffer, pH 7.4 (Electron Microscopy Sciences 16536-06) for 10 min. Sciatic nerves were dissected and post-fixed in perfusion solution overnight at 4 degrees, followed by embedding in a plastic resin (Winters et al., 2011, Mironova et al., 2018). Semi-thin sections were cut at 0.5 µm thickness and stained with 1% toluidine blue solution. LM images were taken distal to the injury site using a 100 X oil objective on Nikon E600 light microscope equipped with a DS-Fi3 camera and Nikon NIS-Elements software. A representative region of interest (ROI) of approximated 18,000 µm2 was selected in the center of each nerve for quantitative analyses. All myelinated fibers within the ROI were segmented using Neurolucida 360 (MBF Bioscience). To determine fiber diameter, axon diameter, myelination, and G-ratio, the shape-adjusted ellipse (SAE) approach was used for size correction (Bartmeyer et al., 2021).

### Behavioral tests

#### Ladder Walk

Adult (9-12 week) male and female mice were placed on a foot misplacement apparatus (horizontal ladder) (Columbus Instruments, 1028SSR). Rungs are 4 mm wide with a 12 mm spacing and mice were recorded while walking from rung 1-50. Training on the apparatus was done 10 days prior to SNC. Mice were allowed to run the ladder beam 3 times, with 2 minutes spent in the escape box at distal end of the ladder between trials. For training 1-3 mice were allowed to run the beam concurrently. Females were tested before males and the equipment was cleaned between females and males. Recordings from naïve mice were done 3 days prior to SNC. Each mouse was placed at the start of the ladder beam (rung 0) and allowed to run the length of the beam (rung 50) in 4 trials. The runs were recorded using a Canon Vixia HF R500 camcorder. The mice underwent unilateral (left side) sciatic nerve crush. The right side was sham operated and served as control. Mice were subjected to the ladder beam test at 3, 14, 35, and 42 dpc. For behavioral scoring, videos were uploaded to Adobe Premier Pro and each frame analyzed by an investigator blinded to mouse genotype. Scores were recorded as P (plantar grasp) or M (missed steps) for all 50 rungs. P was only counted for proper placement of all toes and a clean skip of the next rung (no dragging of foot). Dragging was calculated by analyzing each M in all frames. A drag was considered if the top of the foot graced the ladder rung. Drags were then totaled and plotted as number of drags per 50 rungs. Time points were treated independently. Student’s t-tests were performed to assess statistical significance using Prism.

#### Toe-spread reflex

For quantification of the toe-spread reflex we followed established procedures (Wilder-Kofie et al., 2011). Briefly, mice were trained to walk on a plexiglass catwalk towards a dark box where food pellets were placed. Mice were recorded walking across the catwalk 4 times for each timepoint. Videos were paused and quantified at a point in time when the injured foot was in gait, allowing for the greatest potential toe-spread. Photographs were taken from the video recordings of the hindpaws at baseline (before nerve crush), and at different post injury time points (from 3 to 56 days post-SNC, and 3 to 56 days post-dSNC), 5-10 photographs per mouse per timepoint. Photographs were analyzed by measuring the distance between tip of the first and fifth toe using Adobe Premier Pro Studio and Image J.

### Sciatic Nerve Recordings of compound action potentials

Adult *Sarm1* WT and KO male mice were subjected to unilateral mid-thigh sciatic nerve crush injury. At 73– 122 days post-injury, mice were killed by CO_2_ inhalation and the sciatic nerves were dissected on the ipsilateral and contralateral side. Nerves were incubated in artificial cerebrospinal fluid (ACSF), saturated with 95% O_2_ / 5% CO_2_ for a minimum of 60 minutes. ACSF contained (mM): 125 NaCl, 1.25 NaH_2_PO_4_, 25 glucose, 25 NaHCO_3_, 2.5 CaCl_2_, 1.3 MgCl_2_, and 2.5 KCl. The nerves were then transferred to a temperature-controlled recording chamber held at 37± 0.4°C. Each end of the nerve was drawn into the tip of a suction pipette electrode. Stimuli (supramaximal at 3 mA, 50 μsec) were applied at the proximal (spinal cord) end and recordings were made at the distal end of the tibial branch. Much of the stimulus artifact was subtracted by using a second recording electrode placed near the primary recording pipette. The ratio of the recording pipette resistance after insertion of the nerve to that of the pipette alone was measured and used to correct for drift in the amplitudes of the CAP (Fernandes et al., 2014; Shrager and Youngman, 2017; Stys et al., 1991). Signals were filtered at 10 kHz, sampled at 50–100 kHz, and fed into a data acquisition system.

### Schwann cell cultures

Naïve or dSNC-operated distal nerves were harvested from WT and *Sarm1-/-* mice and placed in DMEM/F12 medium (Gibco, #11960-044) supplemented with 2% Penicillin/Streptomycin (Gibco, #15140-122) in ice. Under the microscope, using a pir of fine forceps (Dumont #55, FST), nerves were cleaned of the epineurium and surrounding adipose tissue and fibers were teased. Teased fibers were then transferred to digestion medium containing DMEM/F12 medium (Gibco, #11960-044), 0.25% Dispase and 0.05% Collagenase II (Worthington Biochemical) for 4-7 hours at 37°C. Digestion was stopped by adding 1x volume of DMEM/F12 medium (Gibco, #11960-044) supplemented with 10% FBS (Biotechne, #S11150). Tissue was centrifuged for 200g for 10 min and resuspended in DMEM/F12 medium (Gibco, #11960-044) supplemented with 1% Penicillin/Streptomycin (Gibco, #15140-122), 10% FBS (Biotechne, #S11150), and triturated with a fire-polished glass pipette. After centrifugation, cells were resuspended in the same medium, supplemented with 50 ng/mL of heregulin β1 (Prepro Tech, #100-03) and 2 µM of Fosrkolin (Sigma-Aldrich, # F6886) and plated into glass-bottom poly-L-lysine-coated (10 µg/mL Science cell, #0413) 24-well plates (Cellvis, # P24-0-N). Medium was changed after 24h and subsequently every 3 days.

Immunofluorescence staining protocol consisted of fixation in 4% PFA for 10 minutes followed by blocking with 5% normal donkey serum for 30 minutes. Primary antibodies against p75^NTR^ (R&D, #AF1157; 1:500), c-Jun (CST, #9165; 1:200), and Sox10 (R&D, #AF2864, 1:100) were incubated for 2h at room temperature. Secondary antibodies were diluted at 1:1000 and incubated for 2h at room temperature. Round coverslips (Neuvitro, #GG-12) were mounted with Dapi Fluoromount-G (SouthernBiotech, #0100-20).

Quantification of Sox10+, c-Jun+, and p75^NTR^+ cells was performed in four random 20X fields per well, using ImageJ. For nuclear staining (Sox10 and c-Jun), nuclei were masked using the same threshold parameters for all images, and automatically counted. For cytoplasmic staining (p75^NTR^), cells were manually counted. Cell counts were expressed as the proportion of positive cells from all cells in each field (Dapi+ nuclei count).

### DRG neuron cultures

The dorsal spinal column from adult mice was exposed and the identify of lumbar DRGs established by counting vertebras from the hipbone (Sleigh et al., 2016). L3-L5 DRGs were dissected and harvested into L-15 + N2 (Gibco, 17502048) or N1 (Sigma, N6530) supplement on ice. DRGs were rinsed 5 times in L-15 supplemented with Penicillin/Streptomycin (Life Technologies 15140-122) and minced with micro-scissors in growth media (DMEM Ham’s F-12, 10% FBS, 1X N2 or N1 supplement and 16 nM Cytosine arabinoside; Sigma, C1768). DRGs were digested in collagenase type 2 (10 mg/ml, Worthington Biochemical LS004176) in Ca^2+^, Mg^2+^ free PBS at 37°C for 20 min. Ganglia were dissociated by trituration using a fire polished glass pipette, followed by centrifugation (5 minutes, 160 x g) and two rounds of trituration in wash buffer (DMEM Ham’s F-12, Gibco 10565-018; 10% FBS, Atlanta Biologicals; 1% Penicillin/Streptomycin, Life Technologies 15140-122). Cells were plated in growth media at a density of 0.5 DRGs per well in a 24-well plate (flat bottom plates, Costar), coated with poly-L-lysine 0.01% (MW 70,000-150,000) (Sigma P4707). Plates were coated with poly-L-lysine for 45 minutes at 37°C, followed by rinsing in dH20, air dried and coated with 0.2 mg/mL laminin (Sigma L2020). Cells were placed in a humidified incubator at 37°C, 5% CO_2_ for 20 hours.

### DRG neuron and injury-activated Schwann Cell Co-cultures

3 days following dSNC, distal sciatic nerves from WT and *Sarm1-/-* mice were harvested in 2 mL of a 1:1 solution containing 1XPBS and Wash media (DMEM F-12 + Glutamax (Fisher Scientific, 10-565-042), 10% FBS (Neuromics FBS 002), and 1X Penicillin/Streptomycin (Fisher Scientific, 15-140-122) and minced into 0.5 mm pieces. Nerves were centrifuged at 3000 RPM for 3 min at room temperature. Supernatant was aspirated and the pellet was digested in a solution of 4mg/mL collagenase type II (Worthington Biochemical, LS004176) and 2mg/mL dispase (Sigma-Aldrich D4693) at 37°C for 30 minutes with agitation every 15 minutes. Wash media was added to digested cells and centrifuged at 3000 RPM for 3 minutes, twice. Cells were resuspended in DMEM F-12 + Glutamax, 10% FBS, 1X Penicillin/Streptomycin, and 0.2 µM Forskolin (Sigma-Aldrich F6886) and were plated either directly onto poly-l-lysine (Sigma P4707) coated glass coverslips, or onto a bed of DRG neurons.

For DRG co-cultures two different methods were utilized. First, Schwann cell cultures as described above were left *in vitro* for 24 hours. At the 24-hour mark, naïve WT DRG neurons were cultured as described above and plated directly onto the bed of Schwann cells. They were left *in vitro* for another 24 hours then fixed stained and imaged. Second, naïve WT DRG neurons were cultured as described above for 24 hours. At the 24-hour mark WT or *Sarm1-/-* Schwann cells were cultured as described above and added directly to the bed of DRG neurons for another 24 hours. They were then fixed, stained, and imaged.

### Quantification of neurite outgrowth

Primary DRG neuron cultures were fixed in 4% paraformaldehyde for 15 minutes at RT, followed by 2 rinses in PBS. Cells were permeabilized in 0.3% Triton-X100 (Sigma, T8787) in PBS for 5 minutes at RT. Cells were incubated in blocking buffer, 2% FBS (Atlanta Biologicals), 2% heat shock fraction V BSA (Fisher Scientific, BP1600), 0.3% Triton-x-100 in PBS for 1 hour. Cells were incubated with anti-Neurofilament heavy chain (1:100; Aves Lab, NFH) in blocking buffer overnight at 4°C and rinsed 3x in 0.3% triton-x-100 in PBS, 5 minutes each. Donkey anti-chicken Cy3 (1:200, Jackson, 703-165-155) in blocking buffer was added for 45 minutes at room temperature. Cells were rinsed in PBS for 5 minutes. Hoechst 33342 (1:50,000 in PBS; Invitrogen, H3570) was added for 10 minutes at RT, followed by 2 rinses in PBS. Cells were imaged on a Zeiss Axio Observer Z1 fitted with a Zeiss Axiocam 503 mono camera using the EC PlnN 10x objective. Single plane, tile scans were randomly acquired for each well. For quantification of neurite outgrowth, NFH stained cells with neurites ≥ 30 µm were included in the analyses from randomly acquired tile scans using WIS-Neuromath (Rishal et al. 2013).

### Bulk RNA sequencing of DRGs

Bulk RNA-sequencing datasets from naïve and axotomized WT DRGs were generated previously (Kalinski et al. 2020) and used for comparison with *Sarm1-/-* DRG datasets. Briefly, we harvested ganglia from naïve *Sarm1-/-* mice (n = 3) and at 1d (n = 3), d3 (n = 3), and d7 (n = 3) following bilateral SNC. For each data point, 18 ganglia were collected form three mice, pooled, flash frozen and lysed in Trizol solution for RNA extraction (Chandran et al. 2016; Kalinski et al. 2020). Libraries were indexed and sequenced over two lanes using HiSeq4000 (Illumina) with 75 bp paired end reads. Sequence quality for the dataset described was sufficient that no reads were trimmed or filtered before input to the alignment stage. Reads were aligned to the latest Mouse mm10 reference genome (GRCm38.75) using the STAR spliced read aligner (version 2.4.0). Average input read counts were 58.8M per sample (range 53.4M to 66.2M) and average percentage of uniquely aligned reads was 86.3% (range 83.8% to 88.0%). Raw reads were filtered for low expressed genes and normalized by variance stabilization transformation method. Unwanted variation was removed by RUVSeq (1.20.0) with k = 1. Differentially expressed genes were identified using the bioconductor package limma (3.42.2) with FDR < 0.1 and the resulting gene lists were used as input for Ingenuity pathway analysis (Qiagen). Weighted gene co-expression network analysis was conducted using WGCNA R-package (ver 1.69). Soft thresholding power of 18 was used to calculate network adjacency. CutHeight of 0.3 was used to merge similar co-expression modules. Enrichment analysis for gene set was performed with GSEA (ver 2.2.2) using MsigDB (ver 7.0). Normalized enrichment score (NES) was used to assess enrichment of gene sets. The bulk RNA-seq and scRNA-seq data is available online in the Gene Expression Omnibus (GEO) database (GSE252734).

### *Ex vivo* nerve cultures

Nerve ex-vivo cultures were performed similarly as described (PMID: 32754584). Mice were perfused with ice-cold 1X PBS for 5 minutes as described above. Naïve sciatic nerve trunks (∼1 cm in length) of WT and *Sarm1-/-* mice were harvested in ice-cold 1x PBS and divided into 2 segments of ∼5 mm. Segments were transferred into a 48-well plate containing 300 µL of DMEM/F12 medium (Gibco, #11960-044) supplemented with 2% Penicillin/Streptomycin (Gibco, #15140-122), 10% FBS (Biotechne, #S11150) and incubated at 37°C for 7, 10, and 14 days. Medium was partially removed and replaced with fresh medium every 5 days. After the designated time points, nerves were removed from culture, placed onto a thick paper to maintain nerves straight, and fixed for 2h in 4% PFA in ice. Nerves were subsequently washed in 1X PBS and placed in 30% sucrose for overnight. Tissues were frozen in OCT as described above and sectioned at 14 µm thickness. Immunofluorescence staining was performed as described above.

Treatment with ErbB inhibitor Canertinib was performed using the same *in vitro* conditions described before. Canertinib 50 µM or vehicle (4-Methylpyridine) were added to the nerve explants upon start of the cultures. After 3 or 10 days, nerves were processed for histology as described.

### Western blotting

Sciatic nerves were dissected and lysed in radioimmunoprecipitation assay (RIPA) buffer (150 mM NaCl, 50 mM TrisHCl, 1% NP-40, 3.5 mM sodium dodecyl sulfate, 12 mM sodium deoxycholate, pH 8.0) supplemented with 50 mM β-glycerophosphate (Sigma-Aldrich G9422-100G), 1 mM Na_3_VO_4_ (Sigma-Aldrich S6508-10G), and 1:100 protease inhibitor cocktail (Sigma-Aldrich P8340-5ML). Nerves were kept on ice, briefly homogenized with a pestle motor mixer (RPI 299200) and subjected to sonication (Fisher Scientific Sonic Dismembrator Model 500) at 70% amplitude for 3 sec. Lysates were centrifuged at 15,000 rpm at 4°C for 10 minutes (Eppendorf 5424R), supernatant transferred to a new tube, and protein concentration measured with a *DC* Protein Assay Kit (Bio-Rad 5000111) using a photospectrometer at 750 nm (Molecular Devices SpectraMax M5e). Samples (10-20 µg total protein) were diluted with 2x Laemmli sample buffer (Bio-Rad 1610737) containing 5% β-mercaptoethanol (EMD Millipore 6010), boiled for 10 minutes at 100°C, and stored at -80°C for analysis. For SDS-PAGE, equal amounts of total protein were loaded for separation in a 4-20% gel. Proteins were transferred onto PVDF membrane (EMD Millipore IPVH00010) for 2.5 hours at 200 mA in ice-cold transfer buffer (25 mM TrisHCl, 192 mM Glycine, 10% Methanol). Membranes were blocked in 5% blotting-grade blocker (BioRad 1706404) prepared in 1x TBS-T (TBS pH 7.4, containing 0.1% Tween-20) for 1 hour at RT, and probed overnight at 4°C with the following primary antibodies diluted in 1x TBS-T with 3% BSA (Fisher Scientific BP1600): α-SARM1 (BioLegend 696602, 1:250), α-CD11b (Abcam ab133357, 1:1000), α-SCG10 (Novus Biologicals NBP1-49461, 1:1000), α-p75^NTR^ (Neuromics GT15057, 1:1000), α-p75^NTR^ (Cell Signaling #8238, 1:1000), α-c-Jun (Cell Signaling #9165, 1:1000), α-P0 (Aves Lab PZO, 1:2000), α-ERK1/2 (Cell Signaling #9102, 1:1000), and α-MBP (Millipore MAB386, 1:500). Horseradish peroxide (HRP)-conjugated secondary IgGs were obtained from EMD Millipore with the following catalog numbers: AP136P (α-rat), AP182P (α-rabbit), AP106P (α-goat). α-chicken HRP-conjugated secondary Ig antibody was obtained from Aves Lab (H-1004). All HRP-conjugated secondary antibodies were used at half the dilution of the corresponding primary antibody in 3% BSA in 1x TBS-T, and the HRP signal was developed with chemiluminescent substrates from Thermo-Fisher Scientific (Pico Plus 34580 or Femto 34095) or from Li-COR Biosciences (926-95000). Protein band intensity was visualized and quantified in the linear range using LI-COR C-Digit (CDG-001313), Image Studio Software (Version 5.2.5), or ImageJ. For normalization, signals were normalized by the corresponding Erk1/2 signal intensity (loading control) and normalized to proximal site within each mouse or to a common Sham-operated sample for each genotype.

### Analysis of NMJs

Animals were perfused with 1X PBS and 4% PFA as described above, hindlimbs amputated, and post-fixed in 4% PFA/PBS for 20 min. EDL muscles were then harvested and stored in 0.025% Triton-x-100/PBS. For whole-mount staining, EDL muscles were teased using forceps to make smaller sized fibers. Teased muscle fibers were then rinsed once in 1X PBS for 5-10 minutes and incubated in blocking buffer (1% BSA, 5% donkey serum in 0.1% Triton-x-100/PBS) for 1 hour. Muscles were then incubated in primary antibodies diluted in blocking buffer, including mouse anti-beta-Tubulin III (1:200, Sigma, T8578) and chicken anti-neuofilament H (1:1000, Aves Lab NFH) for labeling of nerve fibers, rabbit anti-Synapsin (1:200, Cell Signaling, 5297s) for pre-synaptic terminals, overnight at 4°C while rotating. The next day, muscles were washed three times with 0.1% Triton-x-100/PBS, 30 minutes each and incubated with secondary antibody: donkey anti-rabbit CF®647A at 1:200, donkey anti-mouse CF®488A at 1:200, donkey anti-chicken Alexa 488 at 1:500 for 2 hours at RT. During the last 15-20 min of incubation, α-Bungarotoxin CF®543A was added to samples at 1:75. Longer incubation with α-Bungarotoxin CF®543A results in muscle fiber staining and high background. Muscles were then rinsed twice with 0.1% Triton-x-100/PBS, 30 minutes each and once with 1X PBS for 30 minutes. Stained muscles were mounted on slides and flattened using coverslips and imaged with a scanning Leica SP5 confocal microscope. Each selected field of view (FOV) was imaged at 40x magnification, at 2 µm per Z step up to 60-80 µm.

### Flow Cytometry

Single cell suspensions from injured sciatic nerves were prepared as described (Zhao et al. 2022). Briefly, mice were perfused transcardially with ice cold PBS. Injured nerves were harvested and separated into two regions, a ∼3mm injury site and ∼5mm distal site. Nerve segments from 5 mice were pooled per biological replicate to provide sufficient cell counts for analysis. Nerves were digested in 4 mg/ml collagenase II (Worthington Biochemical, LS004176) and 2 mg/ml dispase (Sigma-Aldrich, D4693), followed by myelin removal using a 30% percoll gradient. The resulting single cell suspension was stained using an established antibody panel to identify major immune cell populations (Kalinski et al. 2020). F4/80-APC (Thermofisher Scientific, 17-4801-80) and Rat IgG2a Isotype Control-APC (Thermofisher Scientific, 17-4321-81) were added to the antibody panel. Data was collected using a FACSCanto II (BD Biosciences) flow cytometer and analyzed using FlowJo 10.8.1 software.

## Acknowledgements

We thank members of the Giger lab for critical reading of the manuscript and Sofia Vitale for excellent technical support. This work was supported by the National Institutes of Health 1R15NS128837 (AK), T32DE007057 (LBS), and R01MH119346 (RG), a Ball State University ASPiRE Junior Faculty Award (AK), and the Dr. Miriam and Sheldon G Adelson Medical Research Foundation (RK, LH, AH, DG, RG).

