## Supplemental Figures for "Sarm1 is not necessary for activation of neuron-intrinsic growth programs yet required for the Schwann cell repair response and peripheral nerve regeneration"

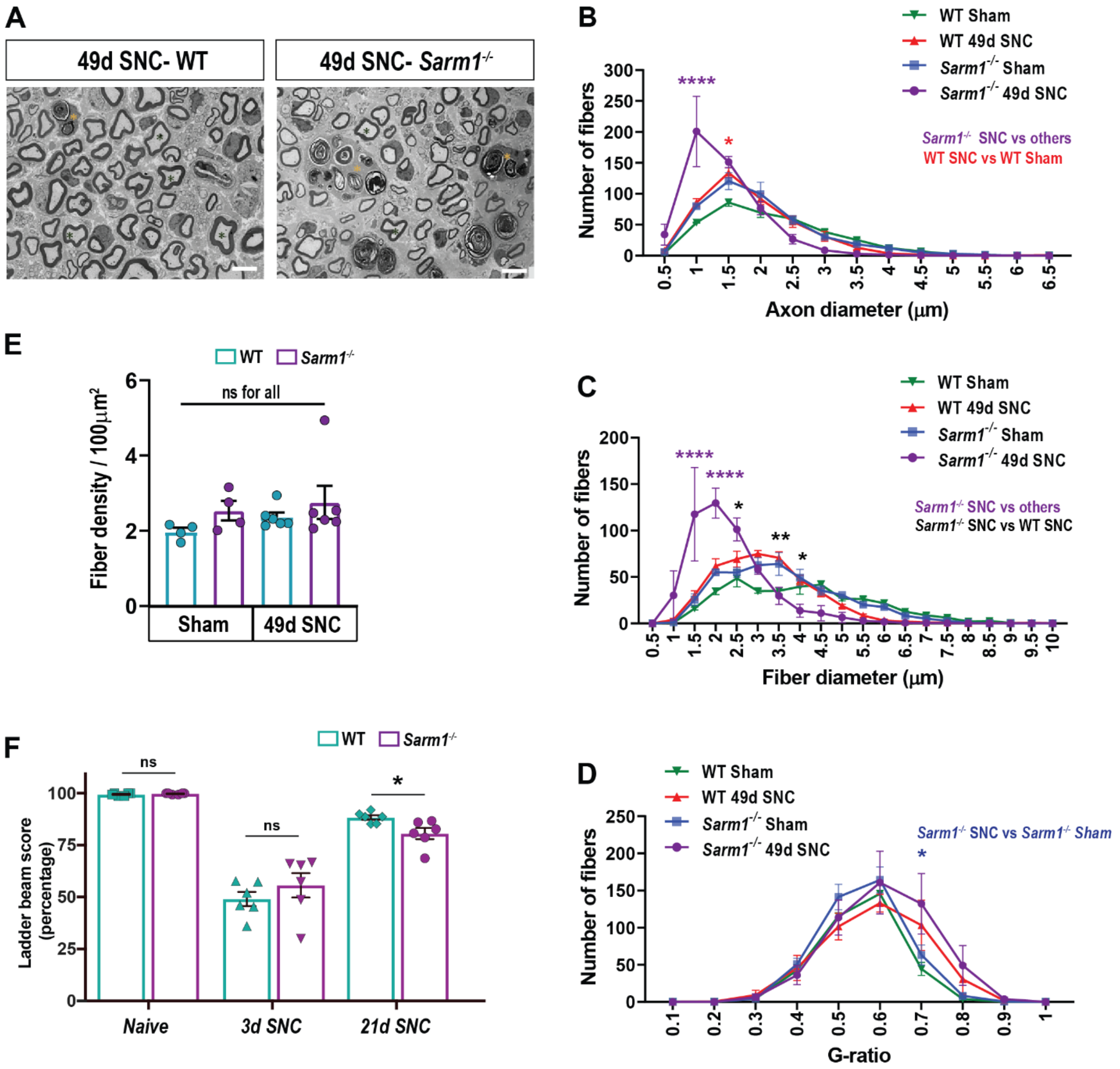


**Figure S1:**

(**A**) Electron microscope (EM) images, axial view, distal to the injury site of WT and *Sarm1-/-* sciatic nerves, 49 days post-SNC. Scale bars, 10 µm.

(**B-D**) Distribution of fibers by axon diameter (**B**), fiber diameter (**C**) and G-ratio (**D**) for WT and *Sarm1-/-* nerves following sham operation and 49 days post-SNC. Quantification of axon diameter, fiber diameter, and G-ratio from EM images, n = 6 injured and 4 sham-operated mice per genotype. Data represented as mean ± SEM. Two-way ANOVA; $^{*}p \leq 0.05$; $^{**}p \leq 0.01$; $^{****}p \leq 0.0001$. Only the most relevant statistical comparisons are shown.

(**E**) Number of fibers per 100 $\mu m^2$ in the sciatic nerve, sham-operated and 49 days post-SNC. N = 6 injured and 4 sham-operated mice per genotype. Data are represented as mean ± SEM. Student's t-test with Mann-Whitney post hoc test; ns, not significant.

(**F**) Horizontal ladder beam score shows the percentage of correct foot placement and foot faults (slips or drags), with 100% being a perfect walking score. *Sarm1-/-* mice show delayed functional recovery at 21 days post-SNC. N = 6 mice per genotype, unilateral sciatic nerve crush (left leg). Data are represented as mean ± SEM. Each data point represents one mouse. $^{*}p = 0.0235$ by two-tailed Student's t-test.

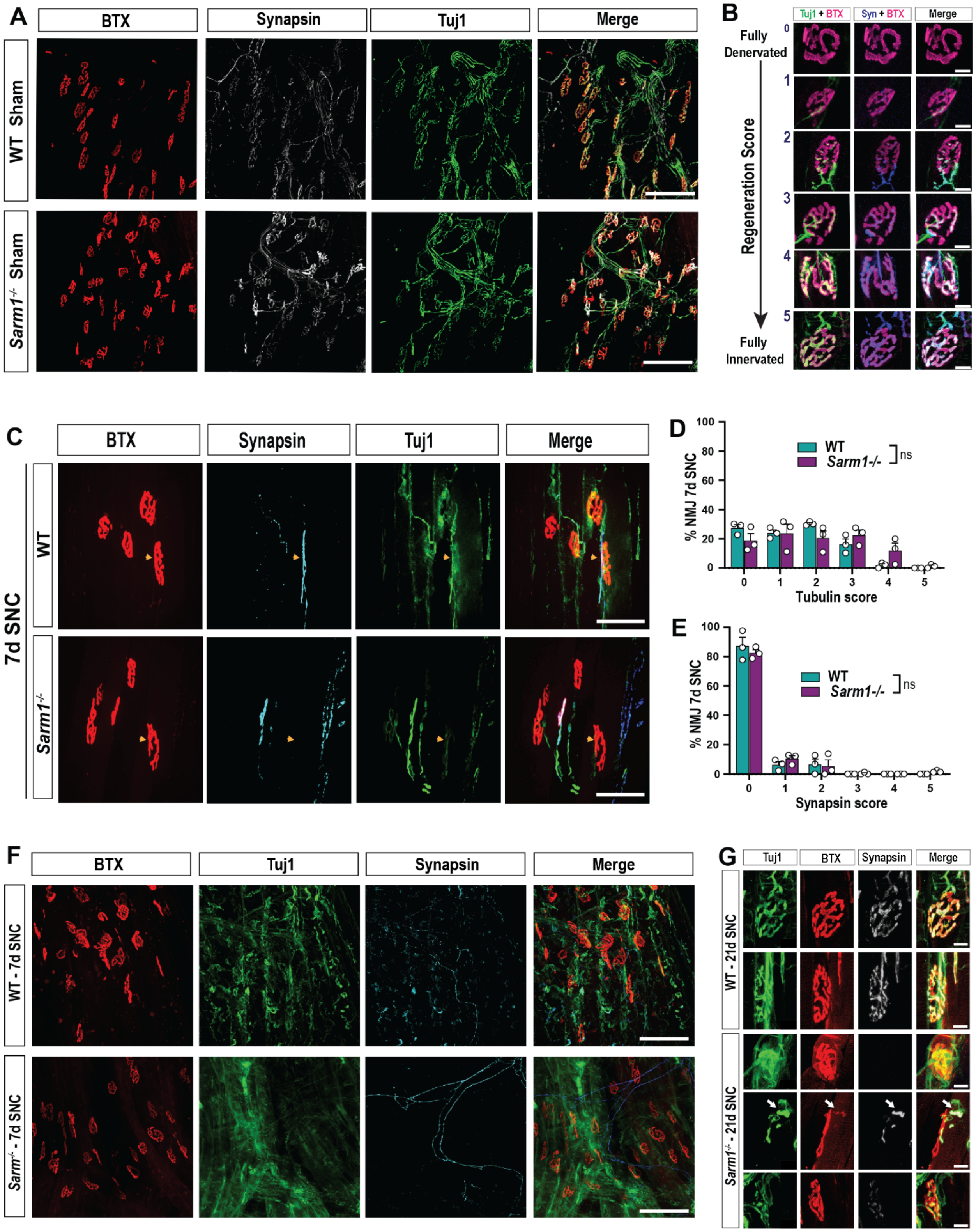


**Figure S2:**

(**A**) Representative whole-mount images of the extensor digitorum longus (EDL) muscles of sham-operated WT and *Sarm1-/-* mice. Neuromuscular junctions (NMJ) stained for postsynaptic endplates (Bungarotoxin, BTX), pre-synaptic terminals (Synapsin), and motor axons (βIII tubulin, Tuj1). Scale bars, 200 μm.

(**B**) Scoring of motoneuron regeneration is based on Tuj1, synapsin, and BTX staining of the neuromuscular junction of WT and *Sarm1-/-* EDL muscles. Completely denervated (score 0); motor axon is approaching, but not innervating an endplate (score 1); >50% of the endplate area is innervated (score 2); <50% of the endplate area is innervated (score 3); motor axon extends throughout the endplate but does not fill the entire space (score 4); and completely reinnervated (score 5). Scale bars, 25 µm.
(**C**) Representative whole-mount images of EDL muscles of WT and *Sarm1-/-* mice at 7 days post-SNC. Neuromuscular junctions (NMJ) stained for postsynaptic endplates (Bungarotoxin, BTX), pre-synaptic terminals (Synapsin), and motor axons (Tuj1). Scale bars, 50 µm.
(**D and E**) Tubulin (**D**) and synapsin (**E**) scores of NMJs quantified at 7 days post-SNC. Data are represented as mean ± SEM. *$p \leq 0.05$; **$p \leq 0.01$; ***$p \leq 0.001$; Student's t-tests; ns, not significant.
(**F**) Whole mount images of EDL muscle 7 days post-SNC in WT and *Sarm1-/-* mice. BTX (red), anti-Synapsin (Cyan or blue) and TuJ1 (green). Scale bars, 200 µm.
(**G**) Representative whole mount images of EDL muscles 21 days post-SNC in WT and *Sarm1-/-* mice. BTX (red), anti-Synapsin (grey) and Tuj1 (green). Arrows show defective target innervation in *Sarm1-/-* mice. Scale bars, 25 µm.

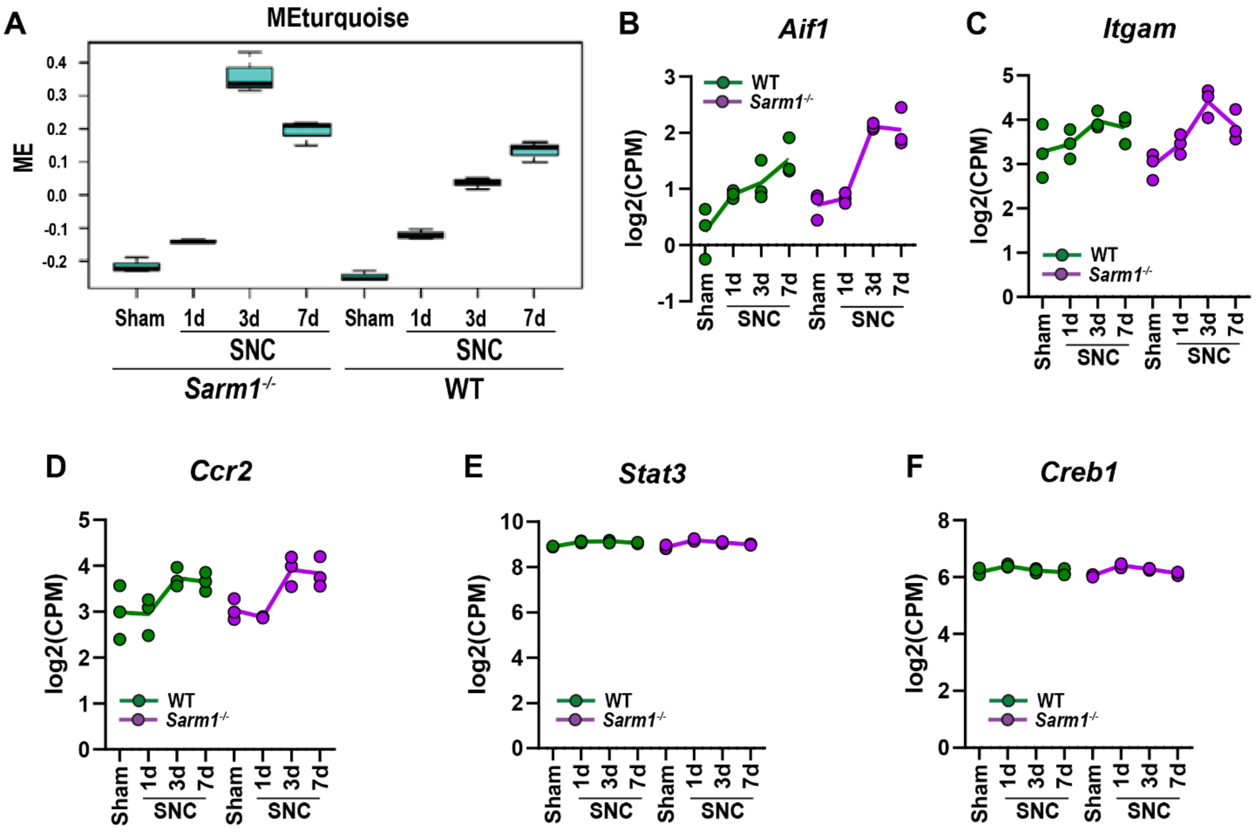


**Figure S4:**

(**A**) Longitudinal analysis of DRG bulk RNAseq datasets. Weighted gene co-expression network analysis (WGCNA) of L3-L5 ganglia harvested from sham-operated, 1, 3, and 7 days post-SNC WT and *Sarm1-/-* mice. Module Eigen (ME) gene expression for the turquoise gene co-expression module, enriched for immune-associated genes, n = 3 mice per genotype and time point.

(**B-F**) Longitudinal expression analysis of gene products in DRGs enriched in macrophages (*Aif1*), innate immune cells (*Itgam/CD11b*), monocytes/macrophages (*Ccr2*) and transcription factors *Stat3* (signal transducer and activator of transcription 3), and *Creb1 (*cAMP responsive element binding protein*)*. Y-axis, log2(counts per million).

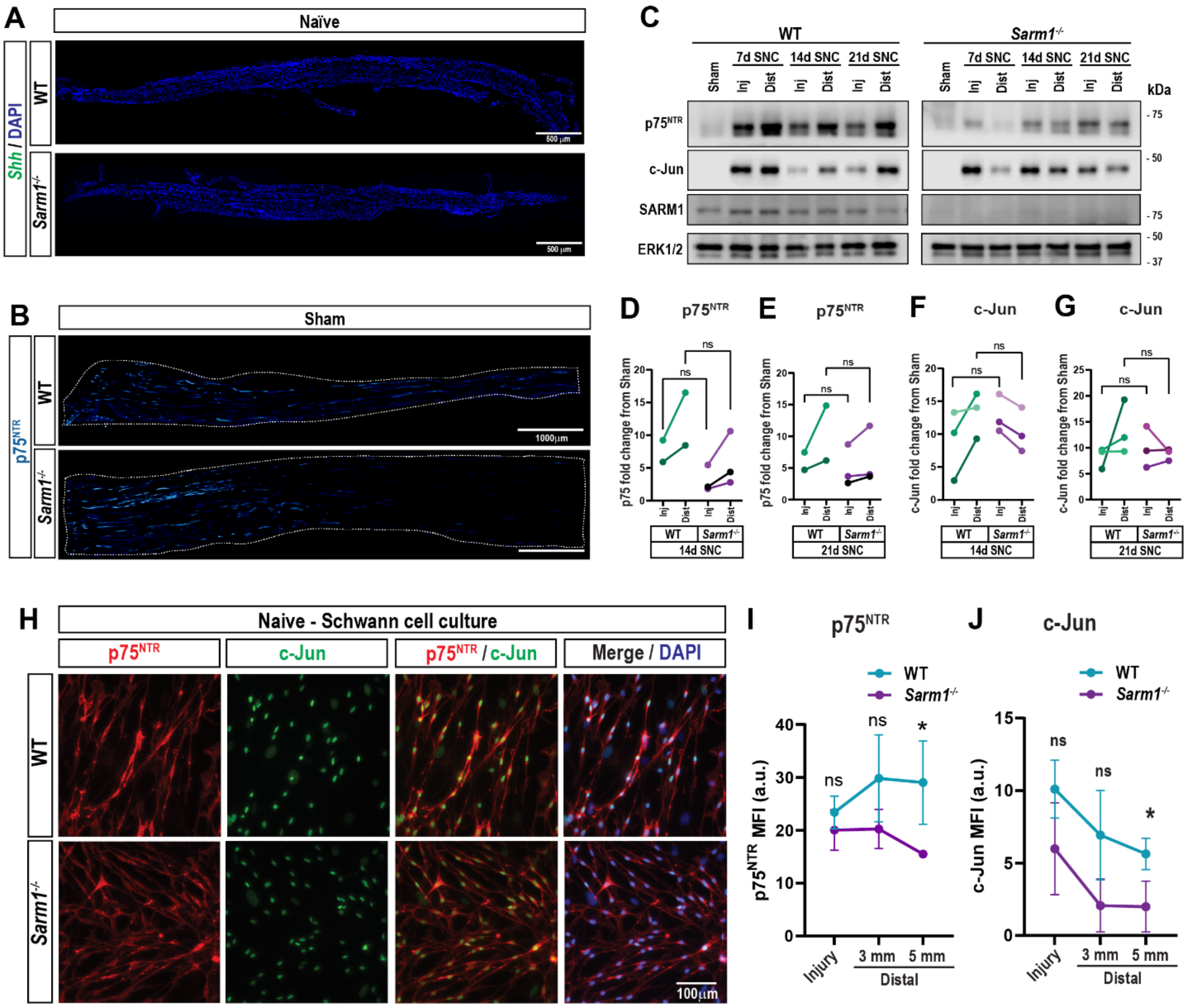


**Figure S5:**
(**A**) Representative images of longitudinal sciatic nerve sections of naïve WT and *Sarm1-/-* mice; RNAscope *in situ* hybridization for *Shh* in green; DAPI in blue. Scale bar, 500 µm; n = 3 mice per genotype.
(**B**) Representative images of longitudinal sciatic nerve sections of sham-operated WT and *Sarm1-/-* mice; stained for p75$^{NTR}$. Scale bars, 500 µm; n = 3 mice per group.
(**C**) Western blotting of sciatic nerve lysates for p75$^{NTR}$, c-Jun, and SARM1 of sham-operated and injured WT and *Sarm1-/-* mice. The injury site and distal nerve were harvested at 7, 14, and 21 days post-SNC and processed separately. ERK1/2 is shown as a loading control.
(**D-G**) Quantification of Western blots shown in (**C**); n = 2-3 biological replicates per genotype, as indicated by the different shades of color. Data were normalized by ERK1/2 and shown as fold-change from sham-operated samples for each replicate (one-way ANOVA; ns, not significant).
(**H**) Primary Schwann cells prepared from naïve WT and *Sarm1-/-* nerves at 7 days *in vitro*, stained for p75$^{NTR}$ and c-Jun. Scale bars, 100 µm; n = 4 mice per genotype.
(**I** and **J**) Analysis of nerves subjected to dSNC; quantification of p75$^{NTR}$ (**I**) and c-Jun (**J**) mean fluorescence intensity (MFI) at the injury site (injury), 3 mm and 5 mm distal to the injury site; n = 3 mice per genotype; one-way ANOVA, *p$\leq$0.05; ns, not significant; a.u., arbitrary units.

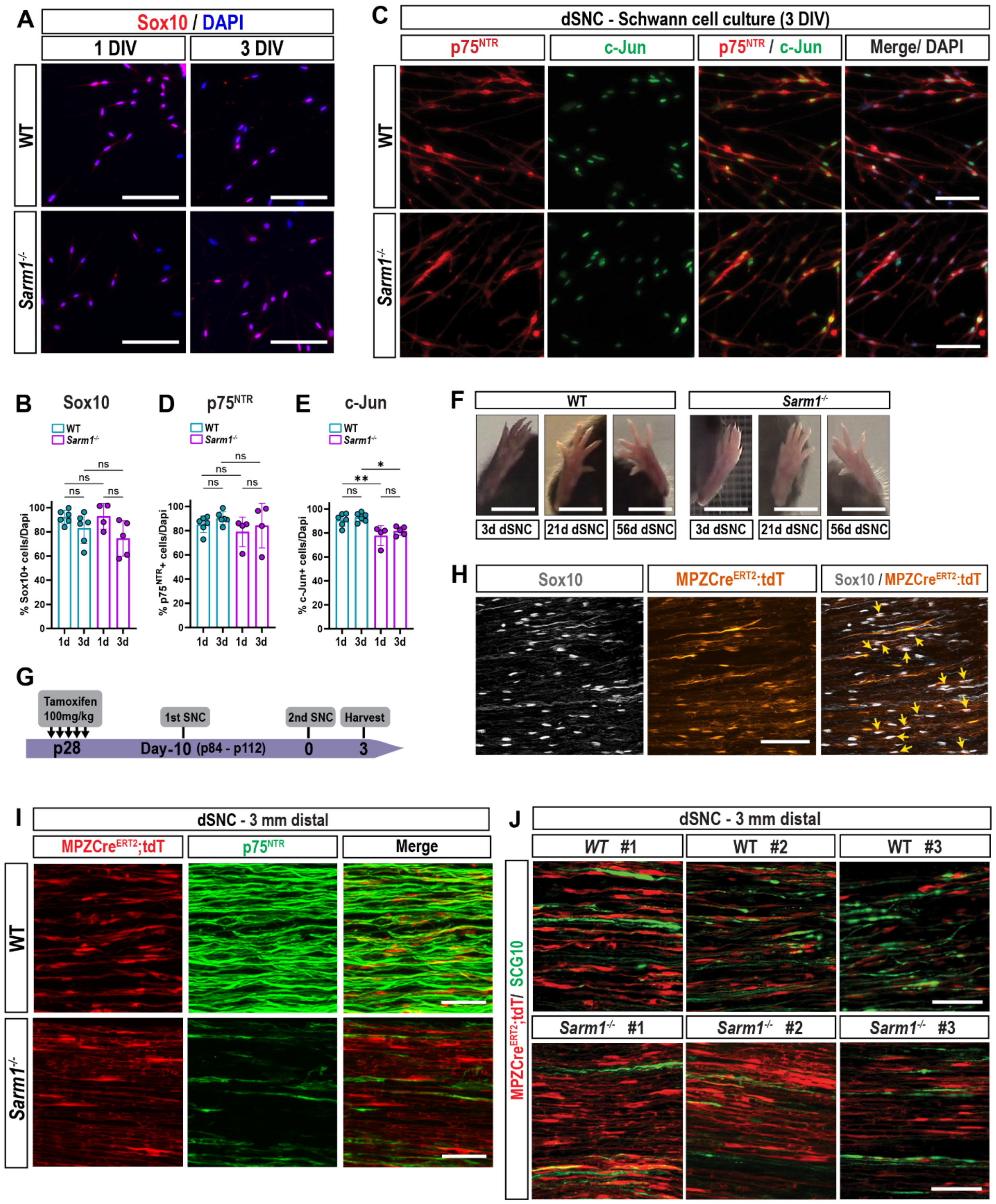


**Figure S6:**
(**A**) Primary Schwann cells at 1 and 3 days *in vitro* (DIV), stained with anti-Sox10 and nuclei (DAPI). Cultures were prepared from the distal nerves of WT and *Sarm1-/-* subjected to dSNC. Scale bars, 100 µm.
(**B**) Quantification of Sox10+ Schwann cells. Y-axis shows the percentage of Sox10+ cells normalized to DAPI+ cells. Data represented as mean ± SEM; n = 4-6 per group. One-way ANOVA; ns, not significant.
(**C**) Primary Schwann cell culture at 3 DIV, stained with anti-p75$^{NTR}$ and anti-c-Jun. Cultures were prepared from the distal nerves of WT and *Sarm1-/-* subjected to dSNC. Scale bars, 100 µm.

(**D-E**) Quantification of p75$^{NTR}$+ (**D**) and c-Jun+ (**E**) Schwann cells. Y-axis shows percentage of stained cells normalized to DAPI+ cells. N = 4-6 mice per group. One-way ANOVA; *p<u><</u>0.05; **p<u><</u>0.01; ns, not significant.
(**F**) Toe-spread reflex, assessed by measuring the distance between the tips of the first and fifth toes in mice subjected to dSNC. Scale bars, 1 cm.
(**G**) Timeline of the dSNC paradigm in WT; or *Sarm1-/-*;*MPZCre$^{ERT2}$;Rosa-tdT* mice.
(**H**) Representative images of longitudinal sciatic nerve sections from naïve *R26$^{tdT/+}$;MPZ$^{CreERT2/+}$* mice stained with anti-Sox10. Yellow arrows show overlapping Sox10 and tdT signals. Scale bars, 100 µm.
(**I**) Representative high magnification images of longitudinal sciatic nerve sections of dSNC WT and *Sarm1-/-*, *R26$^{tdT/+}$;MPZ$^{CreERT2/+}$* mice, stained with anti-p75$^{NTR}$ (green). Shown are SCs (red) 3 mm distally from the injury site. N = 3 mice per genotype. Scale bar, 50 µm.
(**J**) High magnification images of longitudinal sciatic nerve sections of dSNC WT and *Sarm1-/-*, *R26$^{tdT/+}$;MPZ$^{CreERT2/+}$* mice, stained with anti-SCG10 (green). Shown are SCs (red) 3 mm distal to the injury site in WT and mutant mice; n = 3 mice per genotype. Scale bar, 50 µm.

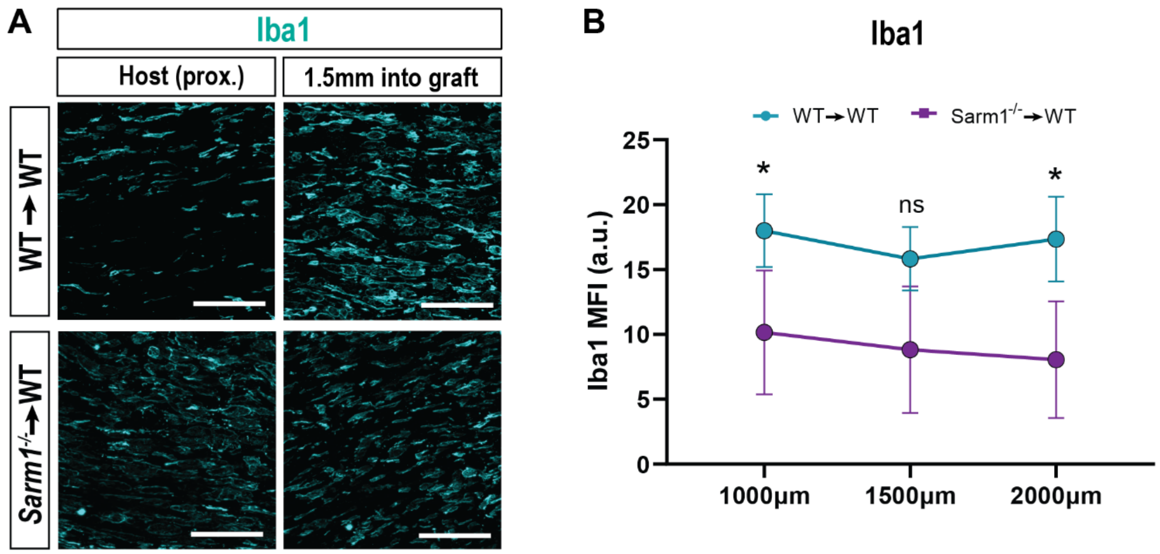


**Figure S9:**

(**A**) High magnification images of WT and *Sarm1-/-* nerve grafts stained with anti-Iba1 14d post grafting. Scale bars, 100 µm.

(**B**) Quantification of Iba1 mean fluorescence intensity ± SEM at 500 µm intervals within grafts, 14d post grafting; n = 5-6 biological replicates per genotype; Students' t-tests; *p≤0.05; ns, not significant.
